# The brain of *Cataglyphis* ants: neuronal organization and visual projections

**DOI:** 10.1101/2020.02.19.954461

**Authors:** Jens Habenstein, Emad Amini, Kornelia Grübel, Basil el Jundi, Wolfgang Rössler

**Author notes:** shared first authorship (Jens Habenstein, Emad Amini). shared senior authorship (Basil el Jundi, Wolfgang Rössler).

## Abstract

*Cataglyphis* ants are known for their outstanding navigational abilities. They return to their inconspicuous nest after far-reaching foraging trips using path integration, and whenever available, learn and memorize visual features of panoramic sceneries. To achieve this, the ants combine directional visual information from celestial cues and panoramic scenes with distance information from an intrinsic odometer. The largely vision-based navigation in *Cataglyphis* requires sophisticated neuronal networks to process the broad repertoire of visual stimuli. Although *Cataglyphis* ants have been subject to many neuroethological studies, little is known about the general neuronal organization of their central brain and the visual pathways beyond major circuits. Here, we provide a comprehensive, three-dimensional neuronal map of synapse-rich neuropils in the brain of *Cataglyphis nodus* including major connecting fiber systems. In addition, we examined neuronal tracts underlying the processing of visual information in more detail. This study revealed a total of 33 brain neuropils and 30 neuronal fiber tracts including six distinct tracts between the optic lobes and the cerebrum. We also discuss the importance of comparative studies on insect brain architecture for a profound understanding of neuronal networks and their function.

## Introduction

Ants of the genus *Cataglyphis* (Foerster 1850) are thermophilic and live in arid zones of Central and North Africa, the Mediterranean, Middle East and in Central Asia. Their natural habitats comprise deserts, steppes and Mediterranean landscapes (Agosti 1990). Due to high surface temperatures and scarce food sources, *Cataglyphis* ants are solitary foragers and do not employ pheromone trails to recruit nestmates (Ruano et al. 2000). The ants predominantly rely on visual cues during far-reaching foraging trips that may span over remarkable distances (reviewed by: Wehner 2003; Buehlmann et al. 2014; Huber and Knaden 2015). Even in environments that lack any panoramic features, *Cataglyphis* foragers find their way back to their nest along an almost straight line using path integration (for reviews see: Ronacher 2008; Wehner 2009). For path integration, the distance and directional information have to be continuously updated (and stored) during outbound trips. In *Cataglyphis*, the distance information is encoded by a stride-integration mechanism in combination with optic-flow perception (Wittlinger et al. 2006; Pfeffer and Wittlinger 2016; Wolf et al. 2018), and the directional information by celestial cues such as the polarization pattern, the position of the sun, and the spectral gradient (Wehner 1997; Wehner and Muller 2006; Lebhardt and Ronacher 2014, 2015). Even though path integration works without any landmark information, *Cataglyphis* ants use the panoramic skyline and visual landmarks of their surroundings, whenever available, to minimize errors (Wehner and Räber 1979; Collett et al. 1992; Wehner et al. 1996; Wehner et al. 2016). The combination of path integration and landmark-guidance generates a robust navigational compass (Knaden and Wehner 2005) and provides a most successful form of navigation. This is likely to be the case in more than 100 *Cataglyphis* species (Agosti 1990) and many other visually guided ants. More recently, *Cataglyphis nodus*, the species in the focus of our present study, was shown to use a magnetic compass for calibrating their visual compasses during naïve learning (exploration) walks that the ants perform upon leaving the nest for the first time and before heading out on first foraging trips (Fleischmann et al. 2018; reviewed by: Grob et al. 2019).

Both path integration and landmark guidance require sophisticated processing and storage of different visual information in the relatively small brain of the ants. Recent studies have mainly focused on two visual pathways – one projects to the central complex (CX), the other to the mushroom bodies (MB) (for reviews see: Grob et al. 2019; Rössler 2019). The CX pathway is highly conserved across insects and has been shown to integrate skylight cues such as polarized light and the sun in the brains of locusts (Homberg et al. 2003; Homberg 2004; Homberg et al. 2011; Heinze 2014), fruit flies (Sancer et al. 2019; Warren et al. 2019), dung beetles (Immonen et al. 2017; el Jundi et al. 2018), monarch butterflies (Heinze and Reppert 2011; Heinze et al. 2013), several bee species (Pfeiffer and Kinoshita 2012; Zeller et al. 2015; Held et al. 2016; Stone et al. 2017) and *Cataglyphis* ants (Schmitt et al. 2016b; Grob et al. 2017).

In all insects investigated so far, sky-compass information is received by photoreceptors in the compound eyes and transmitted from the medulla of the optic lobe (OL) via the anterior optic tract (*AOT*) to the anterior optic tubercle (AOTU). Further neuronal projections connect the AOTU to the lateral complex (LX) and, from there, to the CX. The CX integrates different celestial cues (Heinze and Homberg 2007; Homberg et al. 2011; el Jundi et al. 2014) with other sensory information and represents a high order center for motor control in insects (Strauss 2002; Guo and Ritzmann 2013; Martin et al. 2015; Seelig and Jayaraman 2015). The second pathway to the MB transmits the visual information to be processed in many parallel microcircuits involved in learning and memory formation (for reviews see: Heisenberg 2003; Menzel 2014) as well as multisensory integration of stimuli (Liu et al. 1999; Lin et al. 2007; Kirkhart and Scott 2015). As a special feature in Hymenoptera, projection neurons of the medulla (ME) form a prominent anterior superior optic tract (ASOT) that projects to the collar region of the MB calyces in both hemispheres of ants (Gronenberg 1999, 2001; Ehmer and Gronenberg 2004; Yilmaz et al. 2016; Grob et al. 2017) and bees (Mobbs 1984; Gronenberg 2001; Ehmer and Gronenberg 2002).

Most previous studies focused on these two visual pathways while a comprehensive description of further visual tracts, their target neuropils in the central brain and physiological relevance, are largely missing in Hymenoptera. The reason for this, most likely, is because many of their target regions in the cerebrum, an area termed central adjoining neuropil (CANP), lack clear boundaries between the enclosed individual neuropil regions. Although the neuropils of the CANP have gained less attention, they probably play as much a role in the processing of visual information as the previously established neuropils, such as the AOTU, MB or CX. For a similar reason, and to be able to assign other attributes to distinct brain regions, a consortium of insect neuroanatomists introduced a systematic nomenclature for all neuropils and fiber tracts of the insect brain using *Drosophila* as a model insect (Ito et al. 2014). Further studies created 3D maps of the brains of the monarch butterfly (Heinze and Reppert 2012), the ant *Cardiocondyla obscurior* (Bressan et al. 2015), the dung beetle *Scarabaeus lamarcki* (Immonen et al. 2017), and, more recently, the desert locust *Schistocerca gregaria* (von Hadeln et al. 2018).

In this study, we provide a comprehensive map of synapse-rich neuropils and major connecting fiber tracts including all visual pathways in the brain of the thermophilic ant *Cataglyphis nodus*. By means of immunohistochemical staining and anterograde fluorescent tracing, we were able to define and reconstruct 25 paired and 8 unpaired synapse-rich neuropils and found an overall number of 30 connecting fiber tracts. We further labeled and described six major optical tracts and commissures and their projections into the central brain. This extensive neuronal circuitry demonstrates the complexity of the visual system of *Cataglyphis* ants and emphasizes the importance of further studies to understand the complex processing of visual navigational information in the insect brain.

## Material and methods

### Animals

*Cataglyphis nodus* colonies were collected at Schinias National Park, Greece (38°80’N, 24°01’E) and Strofylia National Park, Greece (38°15’N, 21°37’E) and transferred to Würzburg. The ants were kept under a 12h/12h day/night cycle in a climate chamber with constant temperature (24 °C) and humidity (30 %). The animals had permanent access to water and were fed with honey water (1:2) and dead cockroaches twice per week. For all neuroanatomical studies, *Cataglyphis* workers were randomly chosen from queenless colonies.

### Antibody characterization

To visualize the synaptic neuropils and fiber tracts in *Cataglyphis*, we used a monoclonal antibody to synapsin (SYN-ORF1, mouse@synapsin; kindly provided by E. Buchner and C. Wegener, University of Würzburg, Germany) and fluorescently labeled Alexa Fluor® 488-phalloidin (Invitrogen, Carlsbad, CA, USA; Cat# A12379). The presence of synapsin in presynaptic terminals is highly conserved among invertebrates (Klagges et al. 1996; Hofbauer et al. 2009). The specificity of the antibody has been characterized in *Drosophila* (Klagges et al. 1996) and in the honey bee *Apis mellifera* (Pasch et al. 2011) and its affinity was confirmed in numerous neuroanatomical studies on diverse insect species (e.g. Groh and Rössler 2011; Immonen et al. 2017; von Hadeln et al. 2018) including *Cataglyphis* ants (Stieb et al. 2010; Stieb et al. 2012; Schmitt et al. 2016b; Schmitt et al. 2016a). Phalloidin is a cyclic peptide originating from the fungus *Amanita phalloides*, which binds to filamentous actin, e.g. in dendritic tips (Dancker et al. 1975; Frambach et al. 2004), and axonal processes (Rössler et al. 2002). It has recently been used to study brain structures in *Cataglyphis* (e.g. Stieb et al. 2010; Schmitt et al. 2016b). The combination of fluorescently labeled phalloidin and anti-synapsin increased the contrast of tissue structures and, thus, allowed for a more accurate demarcation of the borders within the CANP.

For additional information about the neuropil boundaries and sub-structures, we used a polyclonal anti-serotonin antibody (rabbit@5-HT; Immunostar, Hudson, WI, USA; Cat# 20080). The specificity of the anti-serotonin antibody was previously tested in dung beetles (Immonen et al. 2017) and its functionality has been demonstrated in diverse insect species (e.g. Hoyer et al. 2005; Dacks et al. 2006; Falibene et al. 2012; Zieger et al. 2013; Watanabe et al. 2014). We used Alexa Fluor® 568-goat@rabbit (Molecular Probes, Eugene, OR, USA; Cat# A11011) and CF633 goat@mouse (Biotium, Hayward, CA, USA; Cat# 20121) as fluorescently-labeled secondary antibodies.

### Immunohistochemistry

Ants were anesthetized on ice before the head was cut off and fixed in dental wax coated dishes. A small window was cut between the compound eyes of the head, and the brain tissue was dissected out and covered in ice-cold ant ringer saline (127 mM NaCl, 7 mM KCl, 2 mM CaCl2, 7.7 M Na2HPO4, 3.8 M KH2PO4, 4 mM TES, and 3.5 mM trehalose; pH 7.0). For whole-mount staining, the brains were fixated in 4 % formaldehyde (FA) in phosphate-buffered saline (PBS) overnight at 4 °C. Brains were washed in PBS (3 × 10 min) on the next day and afterwards treated with PBS containing Triton-X (2 % PBST for 10 min, 0.2 % PBST for 2 × 10 min) to facilitate the penetration of the antibodies. After pre-incubation with 2 % normal goat serum (NGS) in 0.2 % PBST (4°C, overnight), samples were incubated in the primary antibody solution (1:50 anti-synapsin, 2 % NGS in 0.2 % PBST) for 4 days at 4 °C. They were rinsed in PBS (3 × 20 min) followed by an incubation in the primary polyclonal antiserum against 5-HT (1:4000 anti-5-HT, 1 % NGS in 0.2 % PBST) for another 4 days at 4 °C. Subsequently, the brains were incubated in the secondary antibody solution (1:250) and phalloidin (1:200) combined with 1 % NGS in PBS for 3 days at 4 °C. The brains were then washed in PBS (4 × 20 min) and post-fixed overnight in 4 % FA in PBS at 4 °C. After rinsing with PBS, the samples were dehydrated in an ethanol series (30, 50, 70, 90, 95, 100, 100 % for 3-4 min each step), cleared in methyl salicylate (35 min; M-2047; Sigma Aldrich, Steinheim, Germany) and mounted in Permount (Fisher Scientific, Schwerte, Germany; Cat# 15820100).

### Neuronal tracing

All neuronal tracings were done by using combinations of tetramethylrhodamine-biotin dextran (Microruby, 3,000 MW, lysine fixable; Molecular Probes, Eugene, OR, USA, D-7162) and Alexa Fluor 488 dextran (10,000 MW, lysine fixable; Molecular Probes, Eugene, OR, USA, D-22910). To trace the visual and olfactory tracts, the animals were anesthetized on ice, mounted on a holder and the head and antennae were fixed by using dental wax. A square window was cut into the head capsule, and trachea and glands were removed. Subsequently, the brain was rinsed with ant ringer saline before the fluorescent tracer was injected with a glass capillary into the neuropil of interest. To distinguish between tracts that were stained through injections into the medulla (ME) and lobula (LO) of the same optic lobe, different fluorescent tracers were used. In addition, the olfactory tracts were analyzed in a different set of brain samples by injecting Microruby into the antennal lobes (AL). After injection, the injection site was rinsed in ant ringer saline and the head capsule was resealed to avoid dehydration of the brain. For anterograde staining of the antennal nerves, the antennae were cut close to their base and Microruby was added on the antennae stump. In all cases, the animals were stored in humidified chambers in the darkness for 5 hours to allow the dye to be transported. The brains were afterwards dissected and fixated in 4 % FA in PBS overnight at 4 °C. To visualize the neuropils that are innervated by the visual tracts, the corresponding brains were additionally incubated with anti-synapsin antibody as described above. Finally, all brains were rinsed in PBS (5 × 10 min), dehydrated in an ethanol series (30, 50, 70, 90, 95, 100, 100, 100%; each step 10 min), cleared in methyl salicylate (35 min) and embedded in Permount.

### Laser scanning confocal microscopy and image processing

Brain samples were scanned with a confocal laser scanning microscope (either Leica TCS SP2 or Leica TCS SP8, Leica Microsystems AG, Wetzlar, Germany) using a 20x water immersion objective (20.0 × 0.7/0.75 NA). The fluorophores were excited with a wavelength of 488 nm for phalloidin and Alexa Fluor 488 dextran, 568 nm for anti-5-HT and Microruby, and 633 nm for anti-synapsin. All samples were scanned at a step size of 4-6 µm in z-direction and at a resolution of 1,024 × 1,024 pixels in xy-direction. Image stacks were processed using ImageJ (ImageJ 1.52p; Wayne Rasband, NIH, Bethesda, MD) and CorelDRAW X8 (Version 20.1.0.708, Corel Corporation, Ottawa, ON, Canada). If necessary, contrast was adjusted in ImageJ. Pairwise 2D-stitching was performed with the stitching plugin in ImageJ (Preibisch et al. 2009).

### 3D reconstruction and labeling

Three-dimensional (3D) reconstructions of neuropils and fiber tracts were based on anti-synapsin, anti-5-HT and phalloidin labeling using the software Amira 2019.1 (FEI, Visualization Sciences Group; Hillsboro, OR, USA; http://thermofisher.com/amira-avizo). In anti-synapsin staining, only synapse-rich regions are stained while most fiber bundles appear as dark regions. On the other hand, phalloidin staining highlights differences of f-actin distributions between fiber tracts and neuropils. The combination of both techniques in whole mount brains enabled us to better visualize the contours of individual fiber tracts and neuropils. Voxels of the structures of interest were manually labeled in two dimensions in the segmentation editor and, afterwards, interpolated using the interpolation tool. The interpolated contours were double checked and manually corrected in case of deviations from the gray values of the image stack. The corresponding 3D models were generated using the SurfaceGen module.

We followed the unified nomenclature for insect brain introduced by Ito et al. (2014) to define known neuropils and fiber tracts. For undescribed structures, we introduced new terms while accurately following the general rules of the nomenclature (Ito et al. 2014). To define the borders of the brain regions of the central adjoining neuropils (CANP), clearly identifiable fiber tracts and neuropils were used as landmarks. In addition, visible structural alterations of the brain tissue and/or serotonergic innervation patterns were used to discriminate between individual neuropils, similar to the technique introduced by Immonen et al. (2017) and Heinze and Reppert (2012). In some cases, no apparent distinction between neuropils was possible. In these cases, the borders were defined based on traceable and reproducible landmark criteria.

The orientation of the *Cataglyphis* brain relative to its body axis is tilted by about 90 degrees in comparison to that of e.g. *Drosophila* (Pereanu et al. 2010; Ito et al. 2014). Consequently, anterior in *Drosophila* appears as dorsal in *Cataglyphis*, posterior as ventral. As the different orientation can be misleading in comparison with other insect brains, this should be kept in mind, especially considering the nomenclature of individual neuropils and their position in the brain (e.g. the superior neuropils are in the anterior brain region in *Cataglyphis*).

## Results

### General layout of the *Cataglyphis* brain

The central brain of Hymenoptera is organized bilaterally and the protocerebrum, deutocerebrum and tritocerebrum are fused into one cerebral ganglion (CRG) with less distinct borders (Dettner and Peters 2011). In *Cataglyphis,* the CRG on its two lateral ends is enclosed by (compared to many other ant species) relatively large optic lobes (OLs) (Figure 1 a-d, f, g). The distal regions of the OLs are directly attached to the compound eye’s retina. The most conspicuous structures of the CRG are the calyces (CAs) of the mushroom bodies (MBs), which are located at the dorsal tip of the brain (Fig. 1 a-c, e-g). The antennal lobes (ALs) contain olfactory glomeruli (Fig. 1 f, g) and extend to the most ventral regions of the GRG. As in all hymenopteran species (Dettner and Peters 2011), the gnathal ganglion (GNG) is fused ventrally to the CRG and demarcates the posterior end of the central brain (Figure 1 a-d, h). In insects, the GNG comprises three neuromeres: the mandibular, maxillary and labial ganglia, which are fused into one brain region (Scholtz and Edgecombe 2006). Several neuropils of the central brain are surrounded by glial cells and, thus, exhibit well-defined boundaries. This applies to the ALs, the neuropils of the central complex (CX), the MBs and the neuropils of the OLs - and is consistent with anatomical studies in other insects (e.g. Brandt et al. 2005; el Jundi et al. 2009; Heinze and Reppert 2012; Ito et al. 2014; Bressan et al. 2015; Immonen et al. 2017; von Hadeln et al. 2018; Adden et al. 2019; Groothuis et al. 2019). The general shape and structural characteristics of these neuropils in the brain of *Cataglyphis nodus* are also similar to the findings in the honey bee (Brandt et al. 2005). The MB and the CX are surrounded by the CANP, which make up a considerable proportion of the *Cataglyphis* brain (Figure1 e-g).

**Figure 1:**
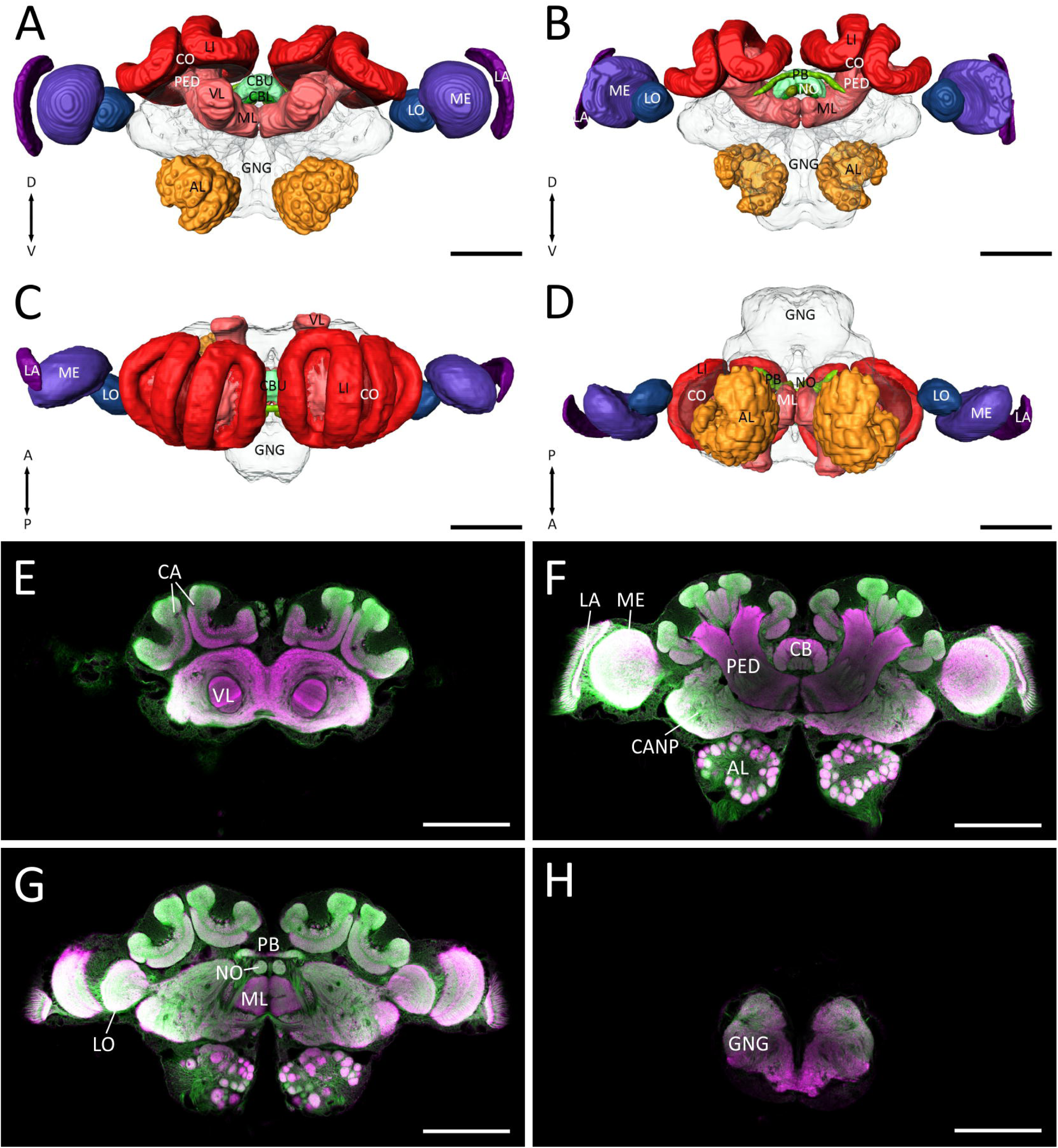
General layout of the *Cataglyphis nodus* brain. **A-D:** Surface reconstructions of the well-defined brain neuropils include the antennal lobes (AL), the mushroom bodies (MB), the optic lobes (OL) and the central complex (CX). The reconstruction of the neuropils is visualized from different views (A: anterior, B: posterior, C: dorsal, D: ventral). The MB can be further subdivided into the calyces (CA), which comprise the collar (CO) and lip (LI), the pedunculus (PED) and the medial and vertical lobe (ML; VL). The CX consists of the central body (CB), the protocerebral bridge (PB) and the noduli (NO) and the optic lobes comprise the lamina (LA), medulla (ME) and lobula (LO). The central adjoining neuropils (CANP, gray) and the gnathal ganglion (GNG, green) are shown as transparent structures. **E-H:** Examples of confocal images from anterior (E) to posterior (H) of the whole brain. The preparation was double-stained with anti-synapsin (magenta) and phalloidin (green) to ensure the highest contrast of the fiber bundles and neuropils. Scale bars = 200 µm.

### Sensory input regions

In contrast to the majority of ant species (Gronenberg 2008), the OLs represent the largest sensory input area in the brain of *Cataglyphis* (Figure 1). They serve as primary processing centers before the visual information is transferred via projection neurons to high order processing and integration sites (reviewed by Rössler 2019). From distal to medial, the OLs consist of the lamina (LA), the medulla (ME) and the lobula (LO) (Figure 2a-c). Based on anti-synapsin and 5-HT staining, the ME can be further subdivided into the inner (iME) and outer medulla (oME). A relatively prominent layer of serotonergic processes demarcates the proximal boundary between the oME and iME (Figure 2d, e). Although tracings indicate that visual information from the dorsal rim area is transmitted via separate paths along dorsal parts of the LA and ME (Grob et al. 2017), no structural features of a distinct dorsal rim area can be localized within the ME or the LA, for example as it has been shown in some other insects (el Jundi et al. 2011; Pfeiffer and Kinoshita 2012; Schmeling et al. 2015; Zeller et al. 2015).

**Figure 2:**
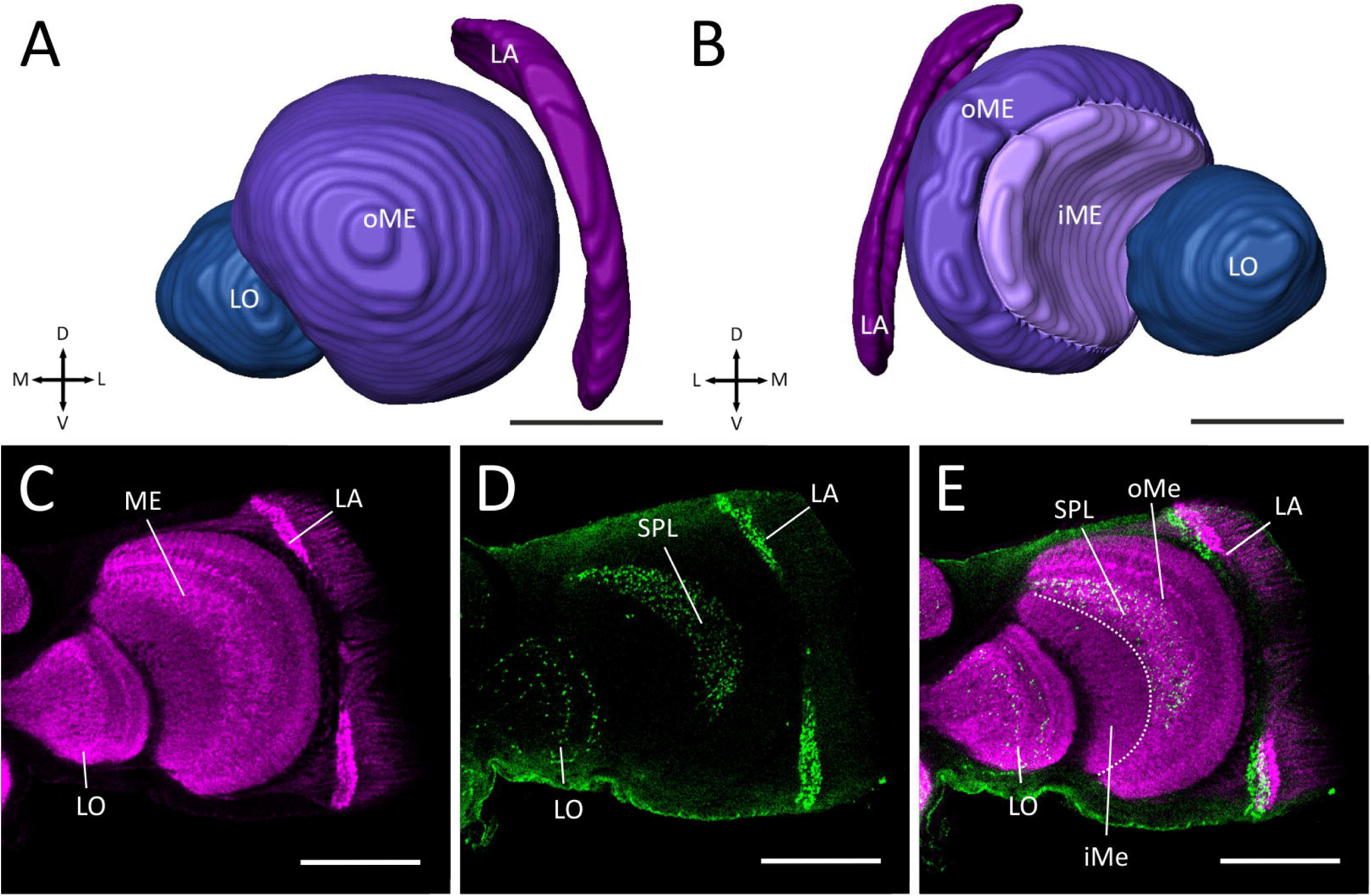
The optic lobes of *Cataglyphis nodus*. **A-B:** Three-dimensional reconstruction of the optic lobes. The optic lobes consist of the lamina (LA), medulla (ME) and lobula (LO). The medulla can be further subdivided into an inner (iME) and outer medulla (oME; A: anterior view; B: posterior view). **C-E:** Confocal images of the optic lobes based on anti-synapsin (C, magenta), anti-serotonin (D, green) labeling, or both (E). The serpentine layer demarcates the border between iME and oME. Scale bars = 100 µm.

The ALs process the olfactory information from olfactory receptor neurons (ORNs) housed in sensilla on the insect antenna. ORN axons project into the glomeruli of the AL. The ALs of *Cataglyphis nodus* contain around 226 olfactory glomeruli (Figure 1a-b, d, f-g) (for glomerulus numbers, see: Stieb et al. 2011).

### High order processing centers

The mushroom bodies are very prominent neuropils in the *Cataglyphis* central brain (Figure 1). The neuroanatomy of the MBs is typical for Hymenoptera: each MB consists of a cup-shaped medial and lateral calyx, a pedunculus as well as a medial and a vertical lobe (Figure 1, 3). The medial and lateral calyces form major input regions of the MB and receive olfactory as well as visual information from afferent projection neurons of the primary sensory neuropils. In *Cataglyphis* each calyx can be subdivided into two subneuropils: the lip (LI) and collar (CO) (Figure 3a, b, d-g). Both LI and CO exhibit characteristic microglomerular structures (Figure 3e-g). Numerous microglomeruli in the visual (LI) and olfactory (CO) subregions of the MB calyx provide thousands of parallel microcircuits forming a rich neuronal substrate for memory formation (Stieb et al. 2012; for reviews see: Rössler 2019; Groh and Rössler 2020). In comparison to many other (mostly olfactory guided) ant species (Gronenberg 1999, 2001), the visual CO is relatively prominent in the *Cataglyphis* MB (Figure 3d-g), further supporting the prominent role of vision. The axons of MB intrinsic neurons (Kenyon cells) innervate the pedunculus and terminate in the medial (ML) and vertical lobe (VL) (Figure 3a, b, d, e-g), which are the major output regions of the MBs (Li and Strausfeld 1997; Strausfeld et al. 2000; Farris 2005; Strausfeld et al. 2009). In *Cataglyphis*, the ML is in the inferior region of the brain, ventral to the CX (Fig. 3f, g), while the VL extends in a perpendicular angle towards the anterior brain surface and forms, together with superior neuropils, the anteriormost structures of the brain (Figure 3a, d, e). On its lateral side, the VL is connected to the calyces by the protocerebral-calycal tract (*PCT*), which represents an intrinsic feedback circuit of the MB (Figure 5a, e). GABAergic *PCT*-neurons have been characterized most extensively in honey bees (Mobbs 1982; Grünewald 1999; Haehnel and Menzel 2010) and may be a general feature of hymenopteran MBs.

**Figure 3:**
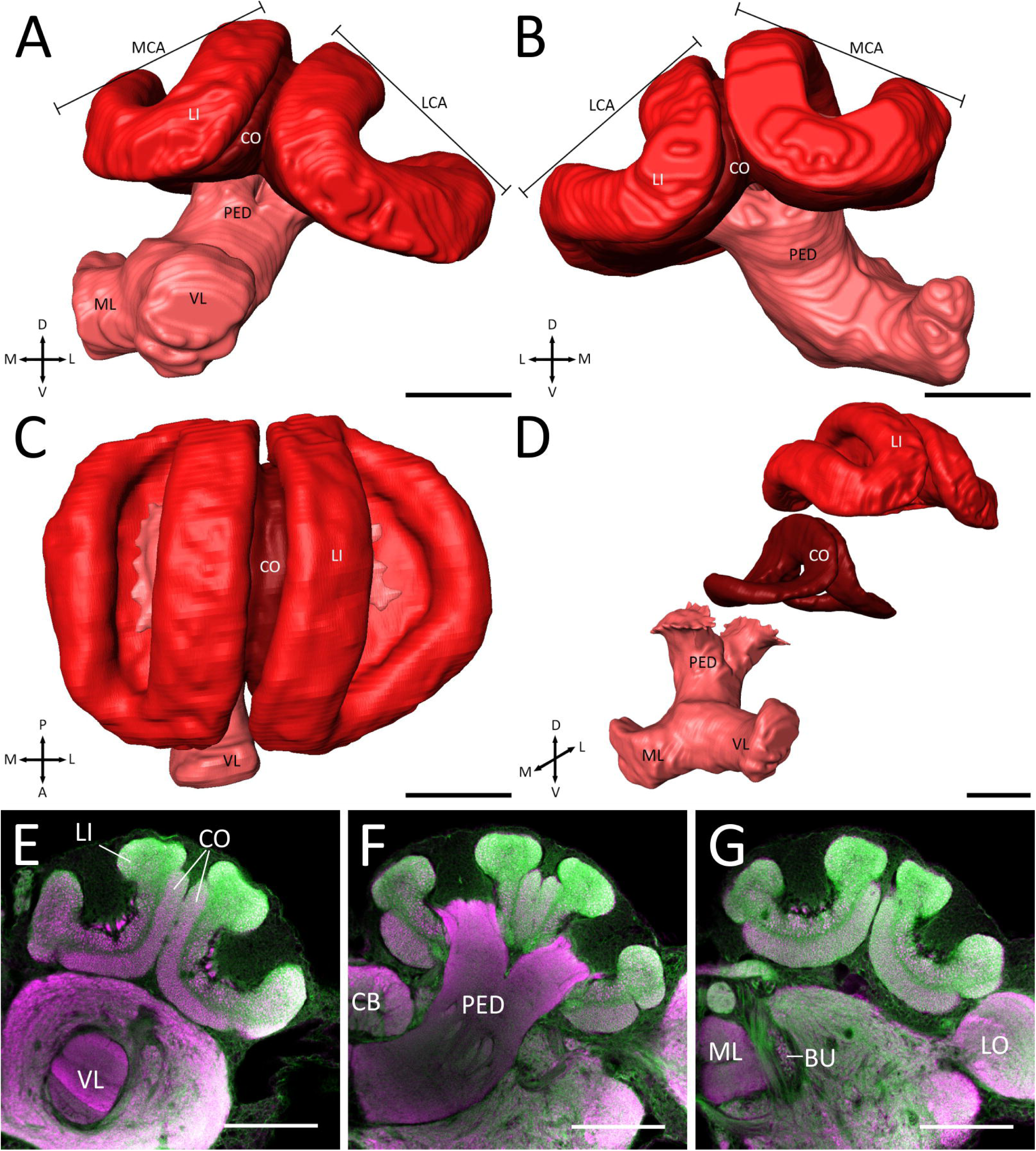
The mushroom body (MB) of *Cataglyphis nodus*. The MB consists of the pedunculus (PED), medial and vertical lobes (ML; VL), and the medial and lateral calyces (MCA; LCA). The calyces can be further subdivided into collar (CO) and lip (LI). **A-D:** Three-dimensional reconstruction of the MB. A: anterior view; B: posterior view; C: dorsal view; D: oblique anteromedial view of the distinct neuropils of the MB. **E-G:** Confocal images of the MB from anterior (E) to posterior (G) of a whole mount preparation (anti-synapsin: magenta; phalloidin: green). BU: bulb; LO: lobula. Scale bars = 100 µm.

Sky compass information converges in the CX before being passed on to premotor centers (Schmitt et al. 2016b; Grob et al. 2017; reviewed by: Rössler 2019). As in other insects, the CX is positioned at the brain midline (Figure 1). It is composed of four distinct neuropils: the upper (CBU) and lower division (CBL) of the central body (CB), the paired noduli (NO) and the protocerebral bridge (PB) (Figure 4). The CBU is often referred to as the fan-shaped body (FB) and the CBL as ellipsoid body (EB). The anteriormost neuropil of the CX is the CBU. To its posterior end, it mounts dorsally on top of the CBL. Both neuropils are located medial to the pedunculi and superior to the MLs of the MBs (Figure 7e-h). More posterior in the brain lies the PB (Figure 7i-k). The only paired structures of the CX in ants are the NO (Figure 4b, d, 7i).

**Figure 4:**
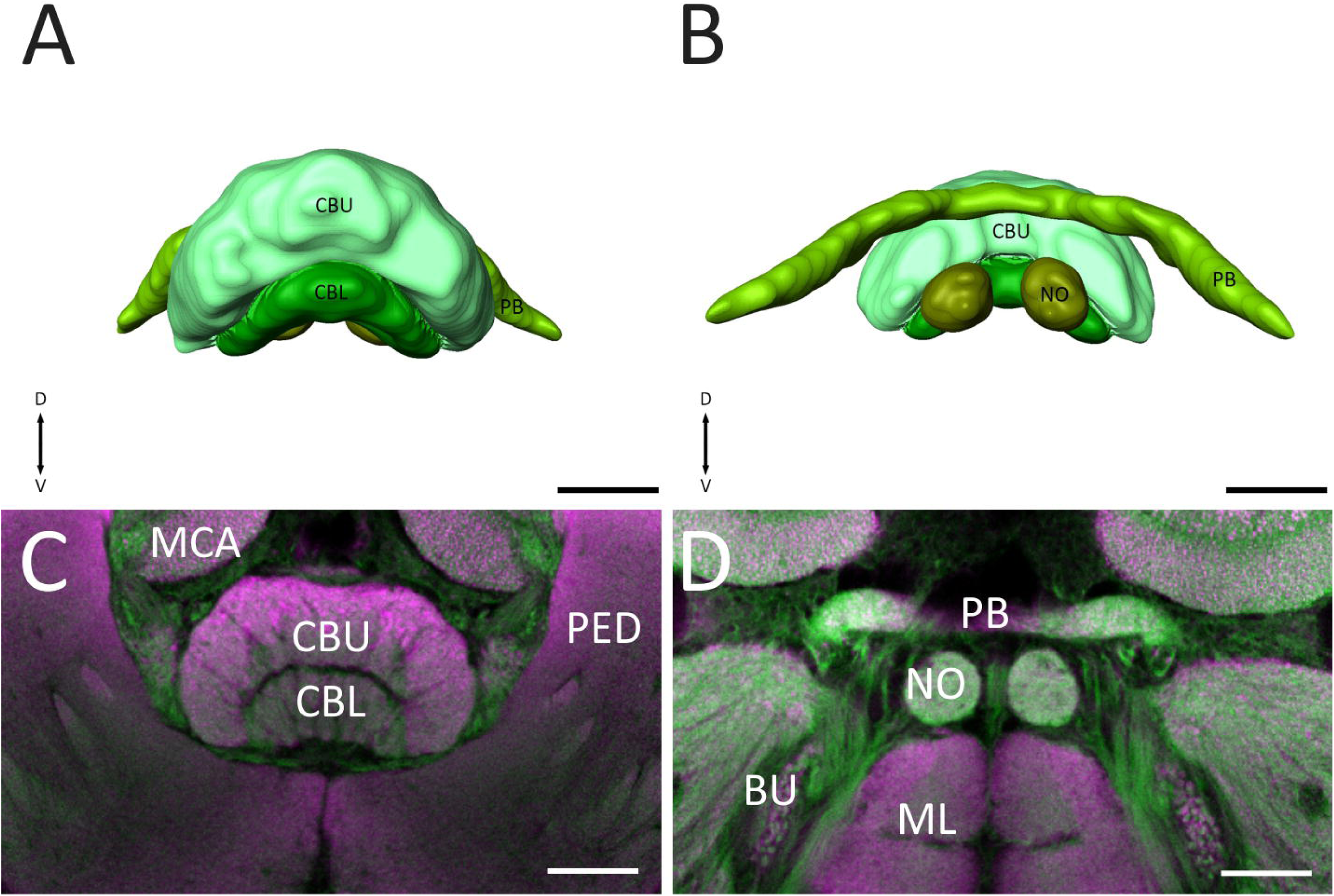
The central complex (CX) of *Cataglyphis nodus*. The CX consists of the upper (CBU) and lower division (CBL) of the central body, the protocerebral bridge (PB) and the paired noduli (NO). **A-B:** Three-dimensional reconstructions of the central complex (A: anterior view; B: posterior view). **C-D:** Confocal images of the CX from anterior (C) to posterior (D) in a whole mount preparation. The preparation was double-stained with anti-synapsin (magenta) and phalloidin (green). BU: bulb; MCA: medial calyx; ML: medial lobe; PED: pedunculus. Scale bars = 50 µm.

### Central adjoining neuropils

In contrast to the major neuropils, the individual regions within the CANP are mostly unsheathed by glial processes and possess less distinct borders in the insect brain (Ito et al. 2014). To define neuropil boundaries, we used prominent fiber bundles as discernable landmarks, as it has also been done in previous work on other insects (e.g. Heinze and Reppert 2012; Immonen et al. 2017; von Hadeln et al. 2018). We first localized and reconstructed the fiber tracts in the *Cataglyphis* brain (Figure 5) to further determine the ambiguous areas of the CANPs. The output pathway of the ALs have been studied in detail, in particular in honey bees and are described as a dual olfactory pathway (Kirschner et al. 2006; reviewed by: Galizia and Rössler 2010; Rössler and Brill 2013). In *Cataglyphis*, we found five output tracts, termed antennal lobe tracts (ALTs), which transfer the olfactory information via projection neurons to higher brain regions: the medial (*m-ALT*), the lateral (*l-ALT*) and three mediolateral tracts (*ml-ALT*; Figure 6a). The *ml-ALT*s are considerably thinner and extend their arborizations into some CANPs: the ventrolateral neuropils, the superior intermediate protocerebrum (SIP) and the lateral horn (LH, Figure 6a, c). In contrast, the *m-ALT* and the *l-ALT* are two of the largest fiber bundles in the entire brain. Both tracts project into the ipsilateral LI and LH (Figure 6a, b). While the *m-ALT* first passes through the LI of the CA (and afterwards into the LH), the *l-ALT* enters the LH first (Figure 6a, b), similar to the findings in the honey bee and in *Camponotus floridanus* (Kirschner et al. 2006; Zube et al. 2008).

**Figure 5:**
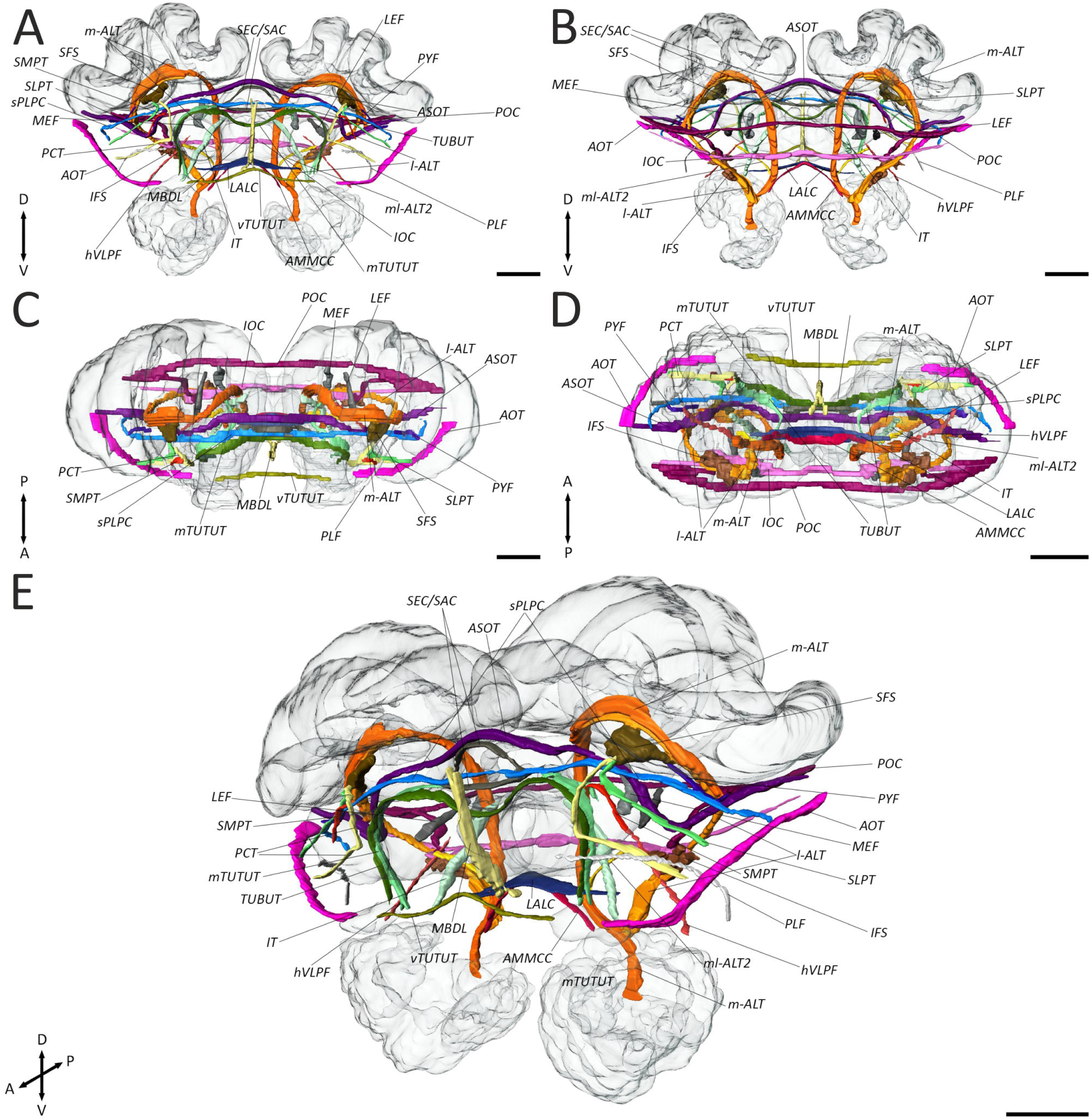
Three-dimensional reconstruction of important fiber bundles in the *Cataglyphis nodus* brain. Only tracts and commissures that could be clearly localized and identified based on anti-synapsin and f-actin phalloidin staining were reconstructed. The mushroom bodies, the central complex and the antennal lobes are shown in transparent. A: anterior view; B: posterior view; C: dorsal view; D: ventral view; E: oblique anterolateral view. *AMMCC*: antennal mechanosensory and motor center commissure; *AOT*: anterior optic tract; *ASOT*: anterior superior optic tract; *hVLPF*: horizontal ventrolateral protocerebrum fascicle; *IFS*: inferior fiber system; *IOC*: inferior optic commissure; *IT*: isthmus tract; *LALC*: lateral accessory lobe commissure; *l-ALT*: lateral antennal lobe tract; *LEF*: lateral equatorial fascicle; *m-ALT*: medial antennal lobe tract; *MBDL*: median bundle; *MEF*: medial equatorial fascicle; *ml-ALT*: mediolateral antennal lobe tract; *mTUTUT* medial tubercle-tubercle tract; *PCT*: protocerebral-calycal tract; *PLF*: posterior lateral fascicle; *POC*: posterior optic commissure; *PYF*: pyriform fascicle; *SEC/SAC* superior ellipsoid/arch commissure; *SFS*: superior fiber system; *SLPT*: superior lateral protocerebrum tract; *SMPT*: superior medial protocerebrum tract; *sPLPC*: superior posterolateral protocerebrum commissure; *TUBUT*: tubercle-bulb tract; *vTUTUT*: ventral tubercle-tubercle tract. Scale bars = 100 µm.

**Figure 6:**
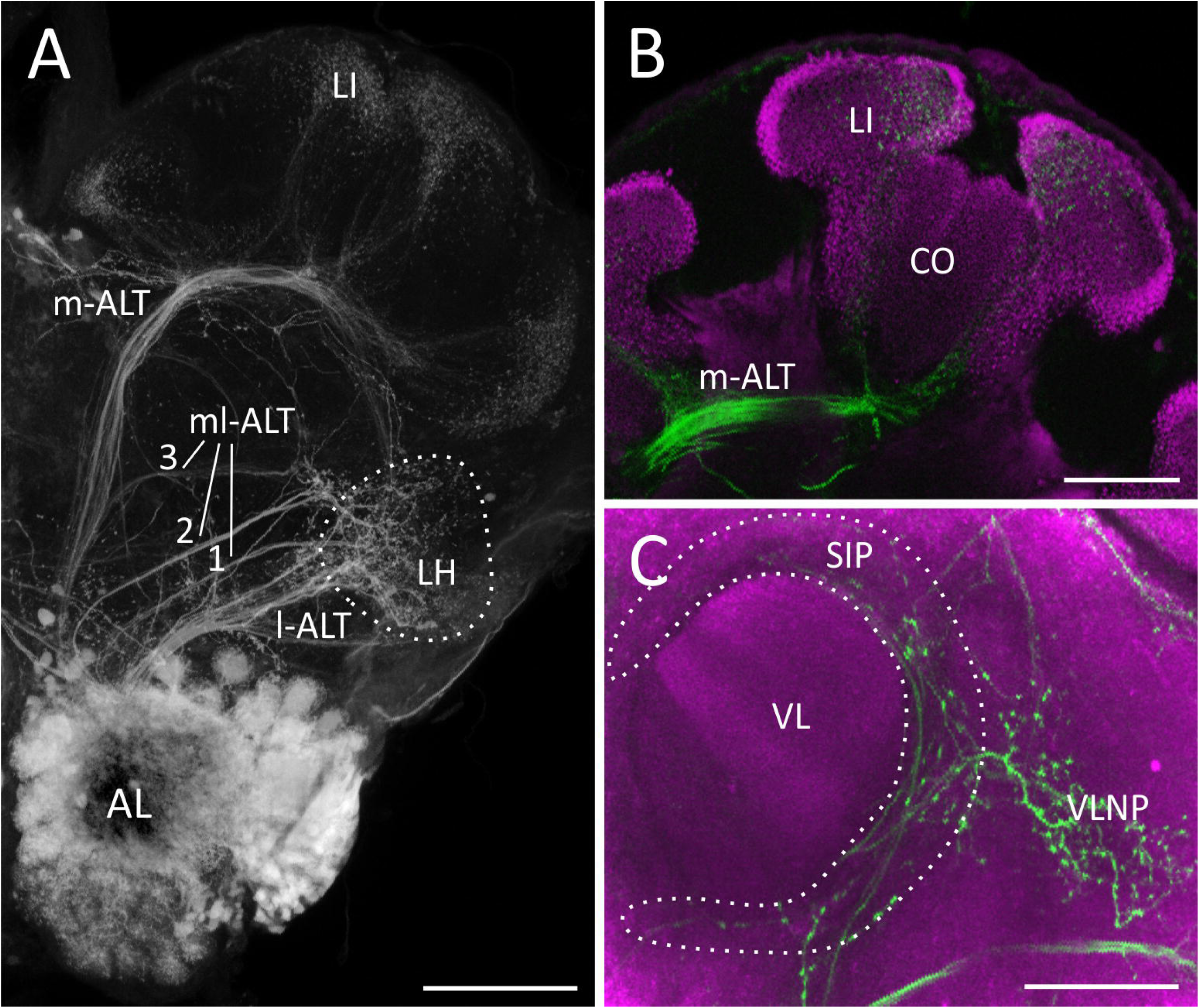
Projections from the antennal lobes (AL) into the central brain. Anterograde staining was obtained by microruby injections (gray/green) into the AL. The pre-synapses are visualized with anti-synapsin labeling (magenta; B,C) **A:** Overview of the antennal lobe tracts. The medial antennal lobe tract (*m-ALT*) and lateral antennal lobe tract (*l-ALT*) project into the lateral horn (LH) and the lip (LI) of the mushroom body (MB). The mediolateral antennal lobe tracts (*ml-ALT*) 1-3 project into different parts of the central adjoining neuropils including the LH. Z-projection from a stack of 27 images with 5 µm step size. **B:** Projections of the *ALTs* into the LI. Z-projection from a stack of seven images with 5 µm step size. **C**: Projections of the *ALTs* into the ventrolateral neuropils and the superior intermediate protocerebrum. Z-projection from a stack of eleven images with 5 µm step size. Scale bars = 100 µm (A) and 50 µm (B,C).

With the help of these tracts as crucial landmarks, the central adjoining neuropils can be subdivided in different brain regions that include the anterior optic tubercle (AOTU), the lateral complex (LX), the superior neuropils (SNPs), the inferior neuropils (INPs), the ventromedial neuropils (VMNPs), the lateral horn (LH), the ventrolateral neuropils (VLNPs) and the periesophageal neuropils (PENPs). Overall, we defined 15 paired and 5 unpaired neuropils in the central adjoining brain region (Figure 7, 8).

**Figure 7:**
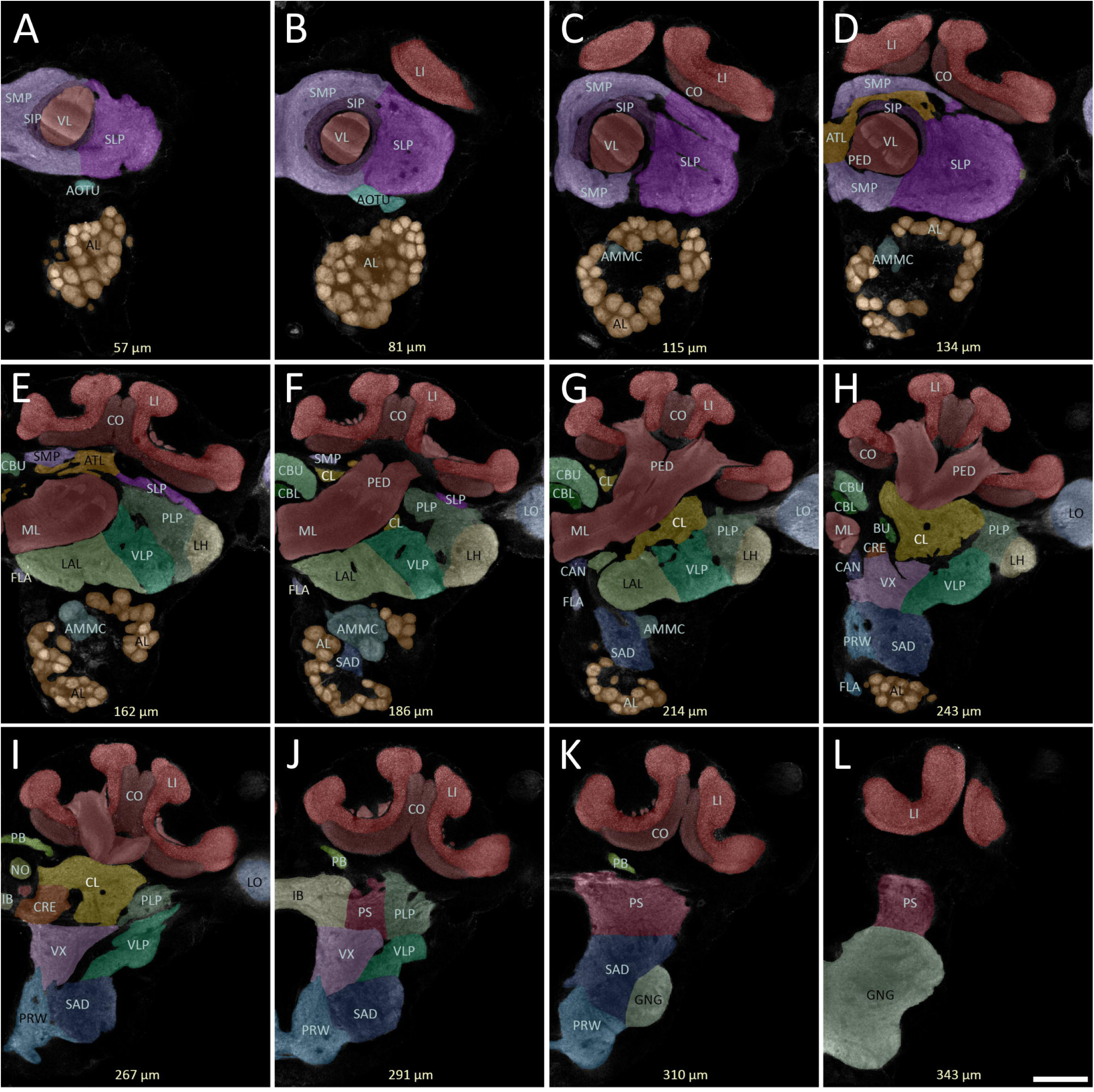
Confocal images (anterior view) of the neuropils of the central brain of *Cataglyphis nodus*. The optical sections (anti-synapsin staining) are shown from anterior (A) to posterior (L). Individual neuropils were labeled with transparent colors to demarcate their boundaries. AL: antennal lobes; AMMC: antennal mechanosensory and motor center; AOTU: anterior optic tubercle; ATL: antler; BU: bulb; CAN: cantle; CBL: central body lower division; CBU: central body upper division; CL: clamp; CO: collar; CRE: crepine; FLA: flange; GNG: gnathal ganglion; IB: inferior bridge; LAL: lateral accessory lobe; LI: lip; LH: lateral horn; LO: lobula; ML: medial lobe; NO: noduli; PB: protocerebral bridge; PED: pedunculus; PLP: posterolateral protocerebrum; PRW: prow; PS: posterior slope; SAD: saddle; SIP: superior intermediate protocerebrum; SLP: superior lateral protocerebrum; SMP: superior medial protocerebrum; VL: vertical lobe; VLP ventrolateral protocerebrum; VX: ventral complex. Scale bars = 100 µm.

**Figure 8:**
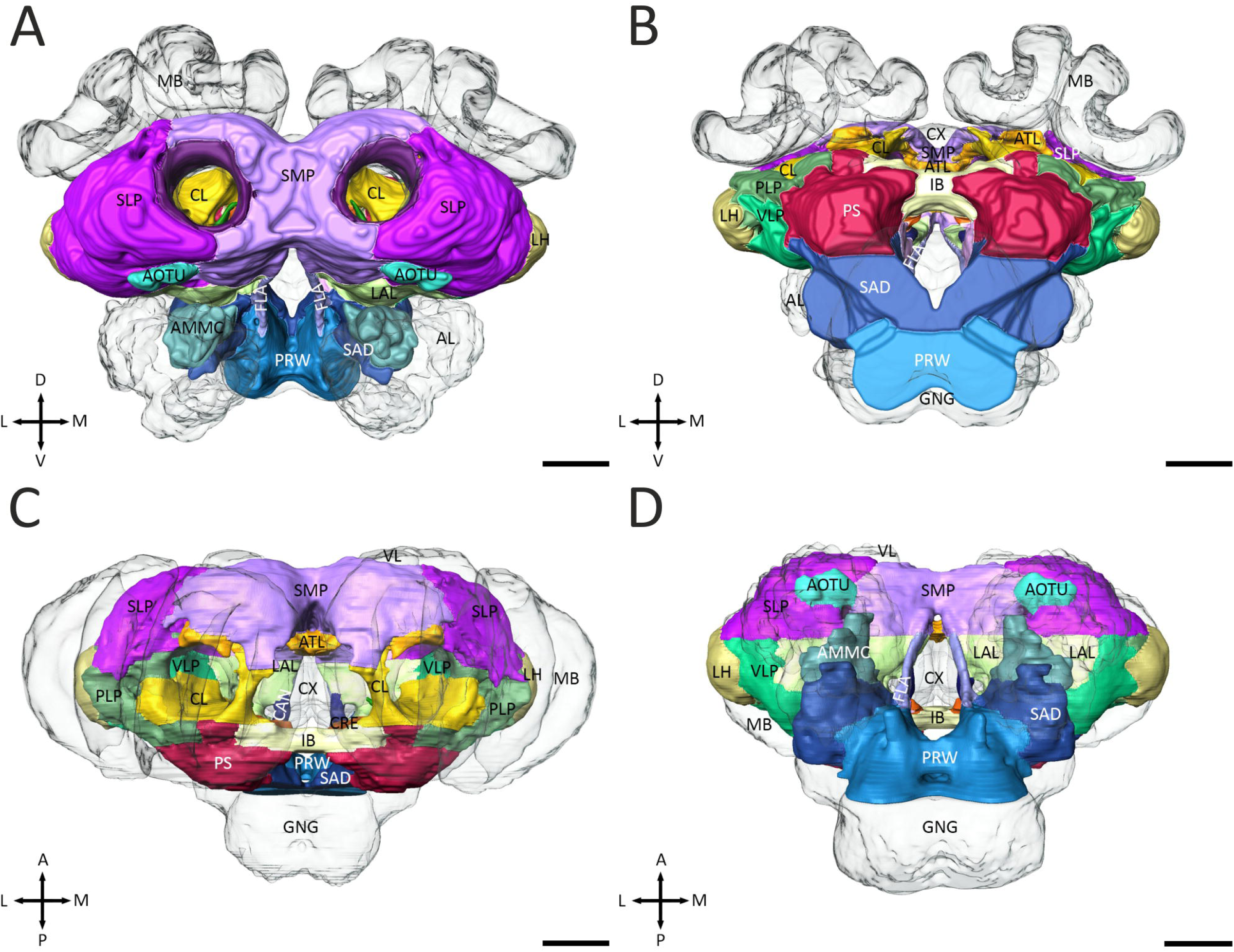
Three-dimensional reconstruction of the central adjoining neuropils. The reconstructions of the neuropils were based on anti-5-HT, anti-synapsin and f-actin phalloidin staining. Antennal lobes (AL), mushroom bodies (MB) and central complex (CX) are shown in transparent. A: anterior view. B: posterior view. C: dorsal view. D: ventral view. AMMC: antennal mechanosensory and motor center; ATL: antler; CAN: cantle; CL: clamp; CRE: crepine; FLA: flange; GNG: gnathal ganglion; IB: inferior bridge; LAL: lateral accessory lobe; LI: lip; LH: lateral horn; PLP: posterolateral protocerebrum; PRW: prow; PS: posterior slope; SAD: saddle; SIP: superior intermediate protocerebrum; SLP: superior lateral protocerebrum; SMP: superior medial protocerebrum; VLP ventrolateral protocerebrum. Scale bars = 100 µm.

### Anterior optic tubercle

The anterior optic tubercle (AOTU) is an important high order visual processing center in insects. As demonstrated in previous studies on locusts, butterflies and honey bees, the AOTU is involved in processing polarized-light information (Pfeiffer et al. 2005; Heinze and Reppert 2011; el Jundi and Homberg 2012) and chromatic cues (Kinoshita et al. 2007; Pfeiffer and Homberg 2007; Mota et al. 2013). In *Cataglyphis*, the AOTU is located superficially in the ventrolateral brain region (Figure 7a, b, 8a) and can be subdivided into two compartments: the upper and the lower subunits (Figure 9a). Projection neurons of the tubercle-bulb tract (*TUBUT*) transmit the visual information from the AOTU to the ipsilateral bulb (BU, Figure 9a). Before the axons finally terminate in the BU, they project around the VL superiorly and pass by the SIP, the antler (ATL) and the clamp (CL). In addition, the AOTUs of both hemispheres are interconnected by the medial (*mTUTUT*) and the ventral tubercle-tubercle tract (*vTUTUT*, Figure 9a). The *mTUTUT* neurons originate from the *TUBUT*. After passing the VL, they bifurcate from the *TUBUT* and cross the midline through the ATL, anteroventral to the fan-shaped body. In contrast to the *mTUTUT* neurons, the *vTUTUT* neurons connect the upper subunits along an almost straight course at the inferiormost border of the superior medial protocerebrum (SMP). Similar intertubercle tracts have been previously described in honey bees (Mota et al. 2011), bumblebees (Pfeiffer and Kinoshita 2012) and locusts (Pfeiffer et al. 2005; el Jundi et al. 2011; el Jundi and Homberg 2012), implying a general characteristic of the interconnection of the AOTUs.

**Figure 9:**
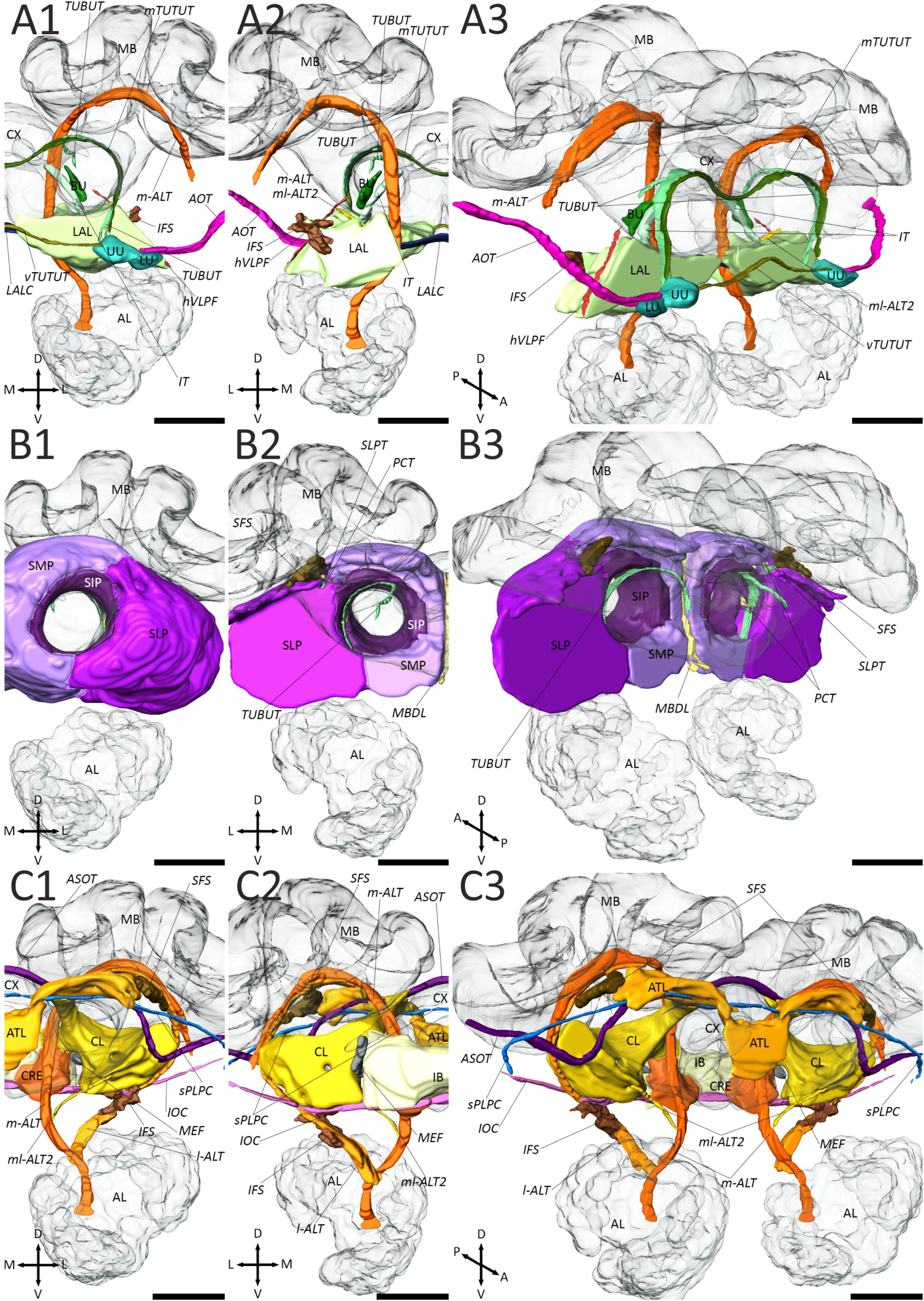
Three-dimensional reconstruction of individual central adjoining neuropils and associated fiber bundles. Antennal lobes (AL), mushroom bodies (MB) and central complex (CX) are shown in transparent. All neuropils are shown from anterior (1), posterior (2), and anterolateral (A3, C3) or posterolateral (B3). **A:** Lateral complex (LX), anterior optic tubercle (AOTU) and associated fiber bundles. The LX can further be divided into the bulb (BU) and lateral accessory lobe (LAL). The AOTU consist of an upper (UU) and lower unit (LU). For the demarcation of the neuropils, the horizontal ventrolateral protocerebrum fascicle (*hVLPF*), the isthmus tract (*IT*), the inferior fiber system (*IFS*), the lateral accessory lobe commissure (*LALC*), the medial antennal lobe tract (*m-ALT*), the medial tubercle-tubercle tract (*mTUTUT*), the mediolateral antennal lobe tract 2 (*ml-ALT 2*), the tubercle-bulb tract (*TUBUT*) and the ventral tubercle-tubercle tract (*vTUTUT*) were used. **B:** Superior neuropils and associated fiber bundles. The superior neuropils consist of the superior intermediate protocerebrum (SIP), the superior lateral protocerebrum (SLP) and the superior medial protocerebrum (SMP). The median bundle (*MBDL*), the protocerebral-calycal tract (*PCT*), the superior lateral protocerebrum tract (*SLPT*) and the *TUBUT* were used as landmarks to define the boundaries of the superior neuropils. **C:** Inferior neuropils and associated fiber bundles. The inferior neuropils comprise the antler (ATL), the clamp (CL), the crepine (CRE) and the inferior bridge (IB). Important fiber bundles for the demarcation of these neuropils are the inferior fiber system (*IFS*), the inferior optic commissure (*IOC*), lateral antennal lobe tract (*l-ALT*), the medial equatorial fascicle (*MEF*), the *ml-ALT 2*, the superior fiber system (*SFS*) and the superior posterolateral protocerebrum (*sPLPC*). Scale bars = 100 µm.

### Lateral complex

The lateral complex (LX) consists of the lateral accessory lobe (LAL) and the bulb (BU) in *Cataglyphis*. The BU is recognizable by its very large microglomerular synaptic structures (Figure 3g, 4d) and presents a prominent synaptic relay of the sky-compass pathway (Homberg et al. 2003; Heinze and Reppert 2011; Pfeiffer and Kinoshita 2012; Held et al. 2016; Schmitt et al. 2016b; Grob et al. 2017; el Jundi et al. 2018). The BU receives sensory information from the AOTU by extrinsic neurons innervating the *TUBUT* (Figure 9a). The *Cataglyphis* brain exhibits only one single bulb per hemisphere which is positioned ventrolateral to the CB (Figure 7h). Its medial border is demarcated by the isthmus tract (*IT*), which connects the LAL with the CB (Figure 9a). The LAL lies inferior to the pedunculus and the CL and more posterior, inferior to the BU (Figure 7e-g). The inferior border is defined by thick glial processes and its medial border by the *m-ALT* (Figure 9a). To its lateral side, the LAL is flanked by the ventrolateral protocerebrum (VLP, Figure 7e-g). The horizontal ventrolateral protocerebrum fascicle (*hVLPF*) and the inferior fiber system (*IFS*) demarcate the boundary between these two neuropils (Figure 9a). As the anterior border between the LAL and the SMP appears to be very ambiguous, the border was set at the level of the anteriormost part of the superior fiber system (*SFS*). The LALs of both hemispheres are interconnected by the lateral accessory lobe commissure (*LALC*, Figure 9a).

### Superior neuropils

The superior lateral protocerebrum (SLP), the superior intermediate protocerebrum (SIP) and the superior medial protocerebrum (SMP) form the superior neuropils (SNP). In *Cataglyphis*, the superior neuropils are the anteriormost region of the brain (Figure 7a-f, 8a, c-d). The largest neuropils of the SNP are the SMP and the SLP. The SLP demarcates the anterolateral border of the brain and extends between the MB calyces and the AL/AOTU all across the anteriormost part of the central brain. To its medial side, the *TUBUT* and the *PCT* define the boundaries of the SMP and the SIP (Figure 9b). The superior lateral protocerebral tract (*SLPT*) outlines the posteroinferior boundary of the SLP to the neighboring posterolateral protocerebrum (PLP, Figure 7e-f, 9b). In contrast to dung beetles (Immonen et al. 2017), we found only one branch of the *SLPT*, which runs from the lateral cell body rind to the *SFS* (Figure 5). The SMP expands to the anteriormost part of the brain across the midline and encloses in a cup-shaped manner on both hemispheres the SIP, the VLs and the anterolateral part of the ATL (Figure 7a-f, 9b). More posteriorly, the median bundle (*MBDL*) and the posteromedial part of the ATL separate the neuropil of both hemispheres (Figure 7d, 9b). Dorsal and anteroventral, respectively, the neuropil is ensheathed by a thick layer of glial cells (Figure 7a-d). *TUBUT*, *PCT*, *SFS* and AOTU demarcate the lateral borders of the SMP (Figure 7a, b, 9b). While SMP and SLP occupy relatively large areas in the brain, the SIP is much smaller. It surrounds the VL in the anteriormost region of the brain and is encircled by the SLP, the SMP and, more posteriorly, by the ATL and the anteriormost edge of the PLP (Figure 7a-d, 9b). As this neuropil is interspersed by many efferent neurons of the VL and the LX, it is recognizable by its less homogenous and intense synapsin-immunoreactivity (-ir). In honey bees, this brain region has been termed ring neuropil, which is characterized by the innervation of the projection neurons from the *ml-ALTs* (Abel et al. 2001; Kirschner et al. 2006).

### Inferior neuropils

The inferior neuropils (INP) occupy the medial brain region around the pedunculus and the ML and lie posterior to the superior neuropils and medial to the ventrolateral neuropils (Figure 7d-i, 8). They comprise the brain areas crepine (CRE), clamp (CL), inferior bridge (IB) and antler (ATL). The ATL and is the anteriormost neuropil of the INP. As in dung beetles (Immonen et al. 2017), the ATL extends across the midline up to the *SFS* and the posteromedial edge of the SLP (Figure 7d-f). The shape of the ATL is closely associated with the processes of the superior posterolateral protocerebrum commissure (*sPLPC*, Figure 9c). The posterior end of the ATL is delineated by a clear glial boundary (Figure 7e, f). Its dorsal and ventral borders are enclosed by the SMP and the SIP (Figure 7d). Since the ATL and most of the neighboring neuropils are obviously contiguous, we defined the borders based on slight differences in the anti-synapsin-ir. More posteriorly in the brain are the CRE, CL and IB located. The CL is wrapped around the posterior part of the pedunculus and is flanked by the *m-ALT* and the *l-ALT* (Figure 9c). It extends from the CBU (dorsomedial) up to the dorsal edge of the LAL (anterior), the ventral complex (VX, posterior) and the ventrolateral neuropils (Figure 7f-i). The CL is separated from the neighboring SMP (superior), ATL (anterolateral), CBU (medial), *m-ALT* (medial), BU (medial), CRE (medial), NO (posteromedial) and the lateral MB calyx (dorsolateral, 7f-i) by thick glial processes. In contrast, most boundaries to the ventral and lateral adjoining neuropils are continuous and were marked by the help of several fiber bundles: the anterior superior optic tract (*ASOT*, anterolateral), the *l-ALT* (lateral), the *ml-ALT 2* (anteroventral), the medial equatorial fascicle (*MEF*, posteromedial), the *IFS* and the inferior optic commissure (*IOC*; both ventral, Figure 9c). The CRE wraps around the ventral and lateral sides of the posterior tip of the medial lobes (Figure 7h, i). In *Drosophila*, this neuropil is heavily innervated by MB extrinsic neurons (Tanaka et al. 2008). It is surrounded by the BU and the CL (lateral), the VX (ventral) and the IB (posterior, Figure 7h-j). On its posterior end, the boundaries of CL and CRE to the IB and the posterior slope (PS) appear rather contiguous. We therefore used the posterior tip of the medial lobes to set the posterior end of these neuropils. The posteriormost subregion of the inferior neuropils is the IB. First identified in *Musca domestica* (Strausfeld 1976), the IB is an unpaired neuropil, which is stretched across the midline. Together with the PS and the posterolateral protocerebrum (PLP), the IB forms the dorsalmost part of the posterior cerebrum in *Cataglyphis* (Figure 7j). The IB is laterally separated from the PS by the *MEF*. The ventral boundary of the IB and the VX, in turn, is defined by the IOC, respectively (Figure 7j, 9c).

### Lateral horn

The lateral horn (LH) is one of the most prominent neuropils of the CANP. It receives direct sensory input from all antennal lobe tracts (Figure 6a) and demarcates the lateral border of the cerebrum (Figure 7d-h, 8). The boundaries of the LH were determined based on the innervation by projection neurons traced along the antennal lobe tracts. In addition, the LH can be distinguished by its brighter synapsin-ir in comparison to its adjacent neuropils. Medial to the LH lie the SLP, the PLP and more ventral, the VLP (Figure 7d-h, 10a).

**Figure 10:**
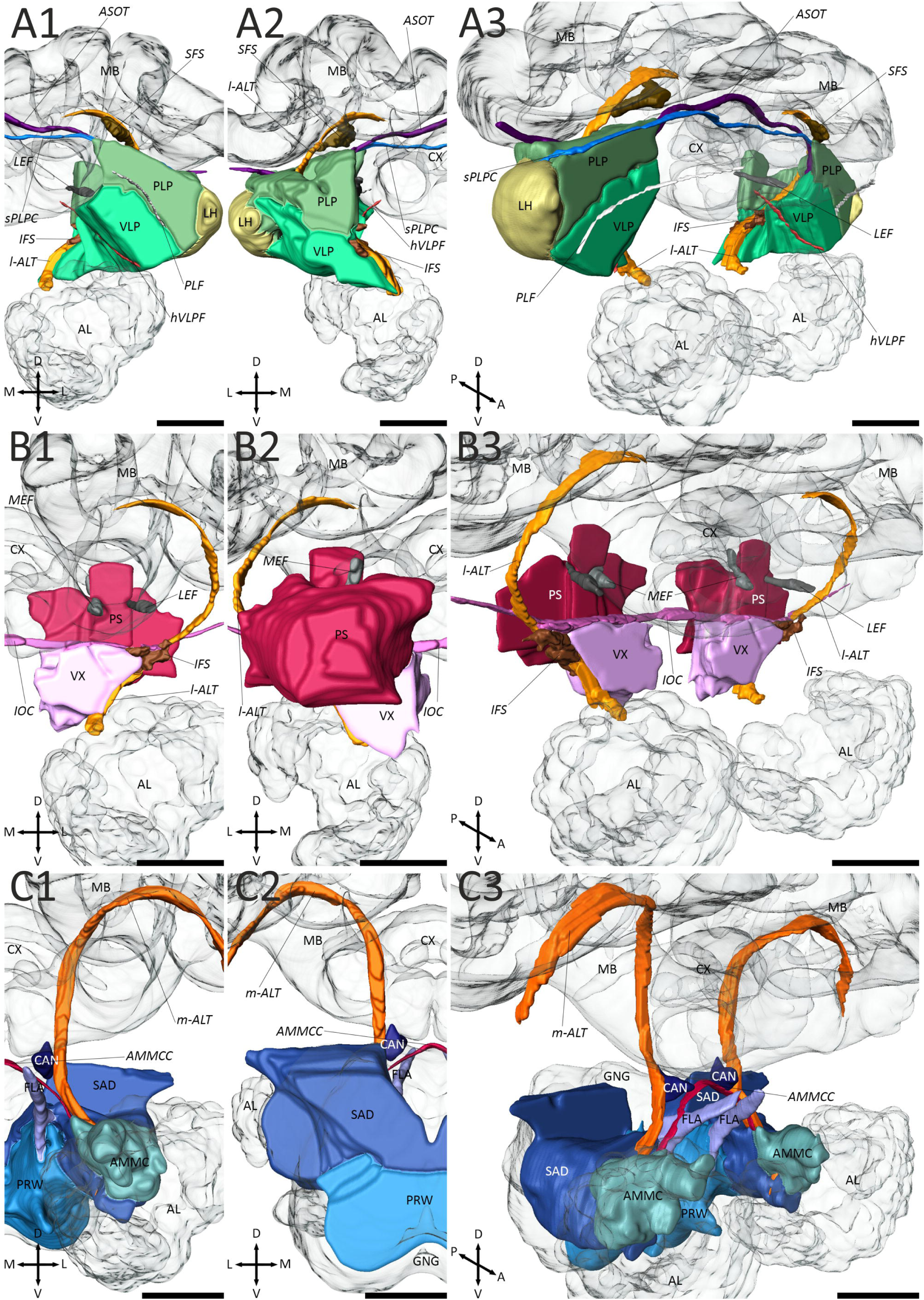
Three-dimensional reconstruction of individual central adjoining neuropil groups and associated fiber bundles (continuing from Figure 9). All neuropil groups are shown in anterior (1), posterior (2) and anterolateral view (3). **A:** Ventrolateral neuropils (VLNP), lateral horn (LH) and associated fiber bundles. The posterolateral protocerebrum and the ventrolateral protocerebrum form the VLNP. The anterior superior optic tract (*ASOT*), the inferior fiber system (*IFS*), the lateral antennal lobe tract (*l-ALT*), the lateral equatorial fascicle (*LEF*), the superior posterolateral protocerebrum commissure (*sPLPC*) and the horizontal ventrolateral protocerebrum fascicle (*hVLPF*) are used to define the boundaries of the neuropils. **B:** Ventromedial neuropils (VMNPs) and associated fiber bundles. The posterior slope (PS) and the ventral complex are the VMNPs in the *Cataglyphis* brain. To demarcate the borders of the VMNPs, the *l-ALT*, the *IFS*, the inferior optic commissure (*IOC*), the *LEF* and the medial equatorial fascicle (*MEF*) were used. **C:** Periesophageal neuropils (PENPs) and associated fiber bundles. The PENPs consist of the antennal mechanosensory and motor center (AMMC), the cantle (CAN), the flange (FLA), the prow (PRW) and the saddle (SAD). Important fiber bundles as landmarks for the demarcation of the PENPs are the medial antennal lobe tract (*m-ALT*) and the antennal mechanosensory and motor center commissure (*AMMCC*). Scale bars = 100 µm.

### Ventrolateral neuropils

The ventrolateral neuropils (VLNP) consist of the ventrolateral protocerebrum (VLP) and the posterolateral protocerebrum (PLP). They are located posterior to the superior neuropils and ventrolateral to the inferior neuropils (Figure 7e-j, 8b-d). The VLNP cover a large area in the *Cataglyphis* brain, expanding from the AL up to the OL and from the anteriormost part of the *SFS* up to the posteriormost regions of the *IFS* and lateral equatorial fascicle (*LEF*, Figure 7e-j, 10a). While the boundaries on the superior and inferior sides are obviously separated by extensive glial sheaths, the medial/lateral and anterior/posterior junctions to neighboring neuropils are very often contiguous (Figure 7e-j). In *Drosophila*, the VLP is divided into an anterior (AVLP) and posterior part (PVLP) based on a more glomerular structure of the PVLP in comparison with the AVLP (Otsuna and Ito 2006; Ito et al. 2014). However, no structural differences could be recognized in this region in *Cataglyphis* and, thus, the neuropil was not further divided into subregions. Unlike in dung beetles (Immonen et al. 2017), the VLP is not characterized by an enriched serotonergic innervation. We therefore defined the borders based on several fiber bundles: the *ASOT* (superior), the posterior lateral fascicle (*PLF*, superolateral), the *l-ALT* (superomedial), the horizontal ventrolateral protocerebrum fascicle (*hVLPF*) and the *IFS* (both medial, Figure 10a). The PLP is demarcated by the *sPLPC* (superior), the *PLF* (inferomedial) the *ASOT* (anteromedial), the *l-ALT* (medial) and the *LEF* (posteromedial, Figure 10a). Lateral to the VLNPs is the LH attached (Figure 7e-h).

### Ventromedial neuropils

Ventral complex (VX) and posterior slope (PS) form the ventromedial neuropils (VMNP) of the *Cataglyphis* brain. They are located posterior to the inferior neuropils and flank the esophagus on both hemispheres of the brain (Figure 7h-l, 8b, c). Because of a lack of unambiguous landmarks, VX and PS were not further divided into subcompartments (namely vest, gorget, and epaulette; superior and inferior PS). The *IFS* is the most important landmark to localize the VX in the brain of *Cataglyphis*. It defines the anterior and posterior end of the neuropil and demarcates, together with the *l-ALT,* the lateral borders to the VLP (Figure 10b). In addition, the *IOC* serves as an apparent dorsal boundary to the neighboring CRE, BU, CL, and more posterior, IB and PS (Figure 7h-j, 10b). In contrast, the ventral boundaries to the prow (PRW) and the saddle (SAD) are rather contiguous but due to a more intense synapsin-ir of the PRW still is clearly recognizable (Figure 7h-j). The PS starts at the level of the IB and is situated at its anteriormost part between IB and PLP, whereas *LEF* and *MEF* form the lateral and medial borders (Figure 7j, 10b). More posterior, the PS expands from the esophagus to the lateral edge of the cerebrum. The SAD and the GNG are localized ventral to the PS (Figure 7k, l).

### Periesophageal neuropils

The periesophageal neuropils (PENP) in the *Cataglyphis* brain comprise the regions of the cantle (CAN), flange (FLA), prow (PRW) and the saddle (SAD), which houses the antennal mechanosensory and motor center (AMMC). These neuropils form the ventralmost region of the brain around the esophagus between the AL and the gnathal ganglion (GNG, Figure 7e-k, 8). The AMMC is the most prominent and probably best studied neuropil of the PENP. It receives primary mechanosensory input from the antennae, providing information about position and movement of the antennae (Homberg et al. 1989; Ehmer and Gronenberg 1997; Ai et al. 2007; Kamikouchi et al. 2009), and gustatory input in a variety of insects (e.g. Jørgensen et al. 2006; Farris 2008; Miyazaki and Ito 2010). To accurately define the borders of the AMMC, in particular to the SAD, we performed antennal nerve backfills (Figure 11). In *Cataglyphis*, the AMMC is adjacent to glomeruli of the AL and more posteriorly it merges smoothly into the SAD (Figure 7c-g). The SAD is a relatively large neuropil and is situated between the LAL and the AL (Figure 7g). More posteriorly, it lies beneath the VX, the VLP and superolateral to the PRW (Figure 7h-j). In its posteriormost region, the neuropil is connected across the midline and its boundaries to the adjacent PS (superior) and the GNG (posterolateral) appear rather contiguous (Figure 7k, 10c). In contrast to the SAD, the CAN is a very small and paired neuropil with clear borders (Figure 7g, h, 10c). In *Cataglyphis*, it fills out a triangular-shaped area in between of the medial lobe (superior), the *m-ALT* (lateral) and the esophagus (medial, Figure 7g, h, 10c). The ventral boundary of the CAN is defined by the AMMC commissure (*AMMCC*) which interconnects the AMMCs of both hemispheres (Figure 10c). Neighboring neuropils are the LAL (lateral), the VX (inferolateral), the SAD (inferolateral) and the FLA (ventral, Figure 7g, h). Unlike in *Drosophila* (Ito et al. 2014), the FLA is heavily surrounded by thick glial processes in *Cataglyphis*. Due to its bar-shaped structure and unambiguously borders it is easily distinguishable from the surrounding neuropils such as the LAL and the SAD (both lateral). It emerges at the root of the median bundle, where it extends lateral from the esophagus and extends up to the anterior tip of the PRW (Figure 7e-h, 8a, d, 10c). The last and most ventral structure of the PENP is the PRW. The structure of this neuropil appears brighter in the anti-synapsin staining than in the remaining adjoining neuropils. Its neighbors are the VX (superior), the FLA (anterior), the SAD (superolateral) and the GNG (posterolateral, Figure 7h-l, 8 a, d).

**Figure 11:**
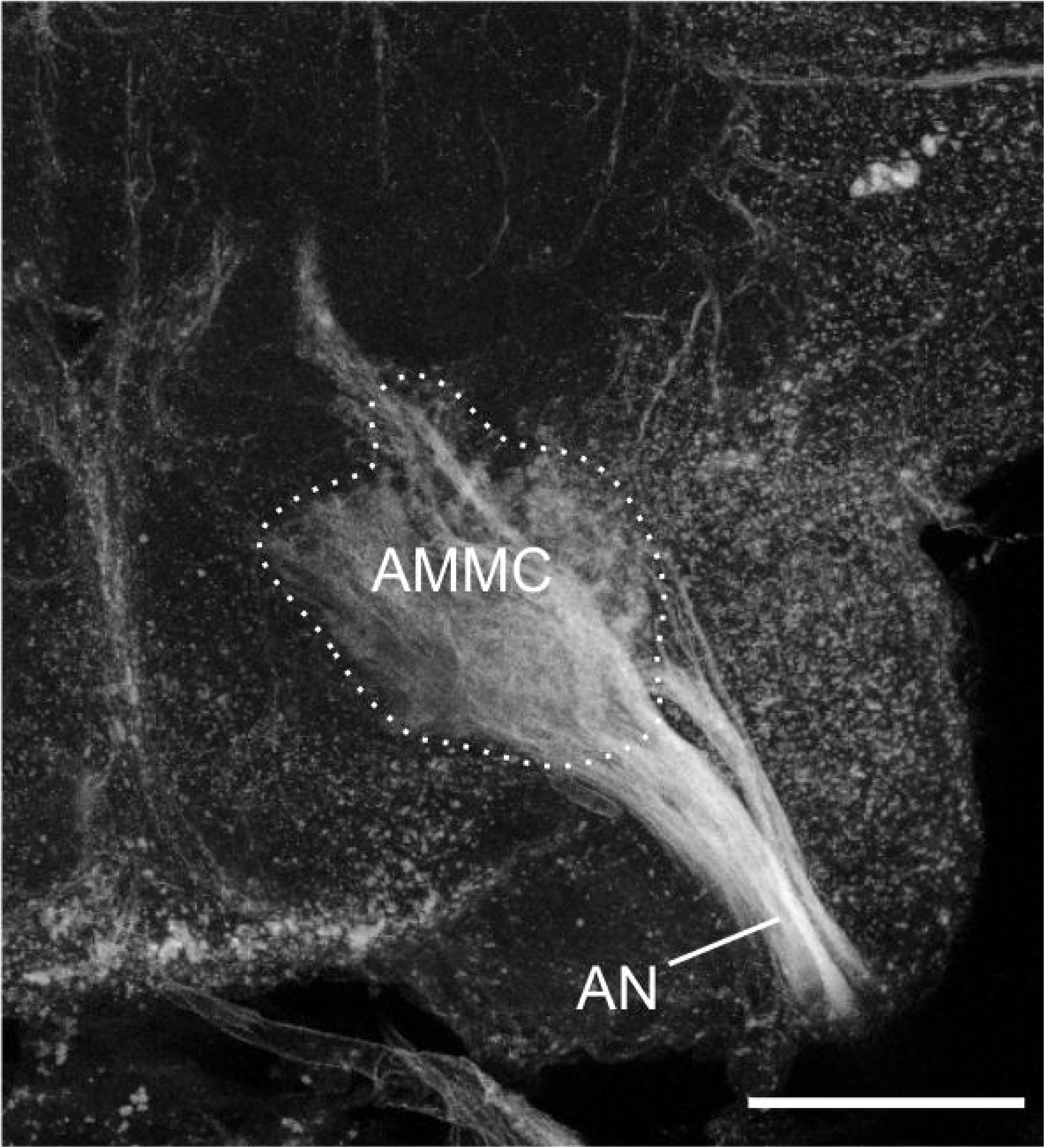
Projection of the antennal nerve (AN) into the antennal mechanosensory and motor center (AMMC). Microruby was used for the anterograde staining. Z-projection from a stack of 21 images with 5 µm step size. Scale bar = 100 µm.

### Visual fibers

To generate an overview of all visual tracts and commissures and their target neuropils, we used neuronal tracer injections into the ME and LO and projected them into the map of synapsin-rich neuropils in the *Cataglyphis* brain. One of the most prominent visual fiber bundles in Hymenoptera is the anterior superior optic tract (*ASOT*) to the lateral and medial calyces of the MBs. In fact, the *ASOT* is also a commissure, which connects the optic lobes of both hemispheres (Figure 12h, 14b). The *ASOT* forms side branches into the ipsilateral and contralateral CO of the MB calyces (Figure 12a-c, h, 13, 14b). The *ASOT* predominantly comprises optic lobe neurons of the ME but also houses a small number of neurons originating from the LO (Figure 12a-c). In contrast to our findings, in honey bees the LO projections run in a separate tract (lobula tract) before they join the *ASOT* (Mobbs 1984; Ehmer and Gronenberg 2002). The *ASOT* can be found anterior in the brain and exhibits a characteristic course through the brain with a sharp bend lateral to the pedunculus, just before it passes the vertical lobes superiorly and crosses the midline dorsal to the central complex (Figure 12a-c, h, 13, 14b). In *Cataglyphis*, the *ASOT* is not the only visual fiber tract with projections into the mushroom bodies. Our tracer injections revealed a second fiber tract of unknown identity, that runs anterior to the *ASOT* and dorsal to the SLP, which we termed optic calycal tract (*OCT*). The *OCT* interconnects the ME and LO with the ipsilateral MB CO but, in contrast to the *ASOT*, does not project into the contralateral brain hemisphere (Figure 12a-c, 13, 14a).

**Figure 12:**
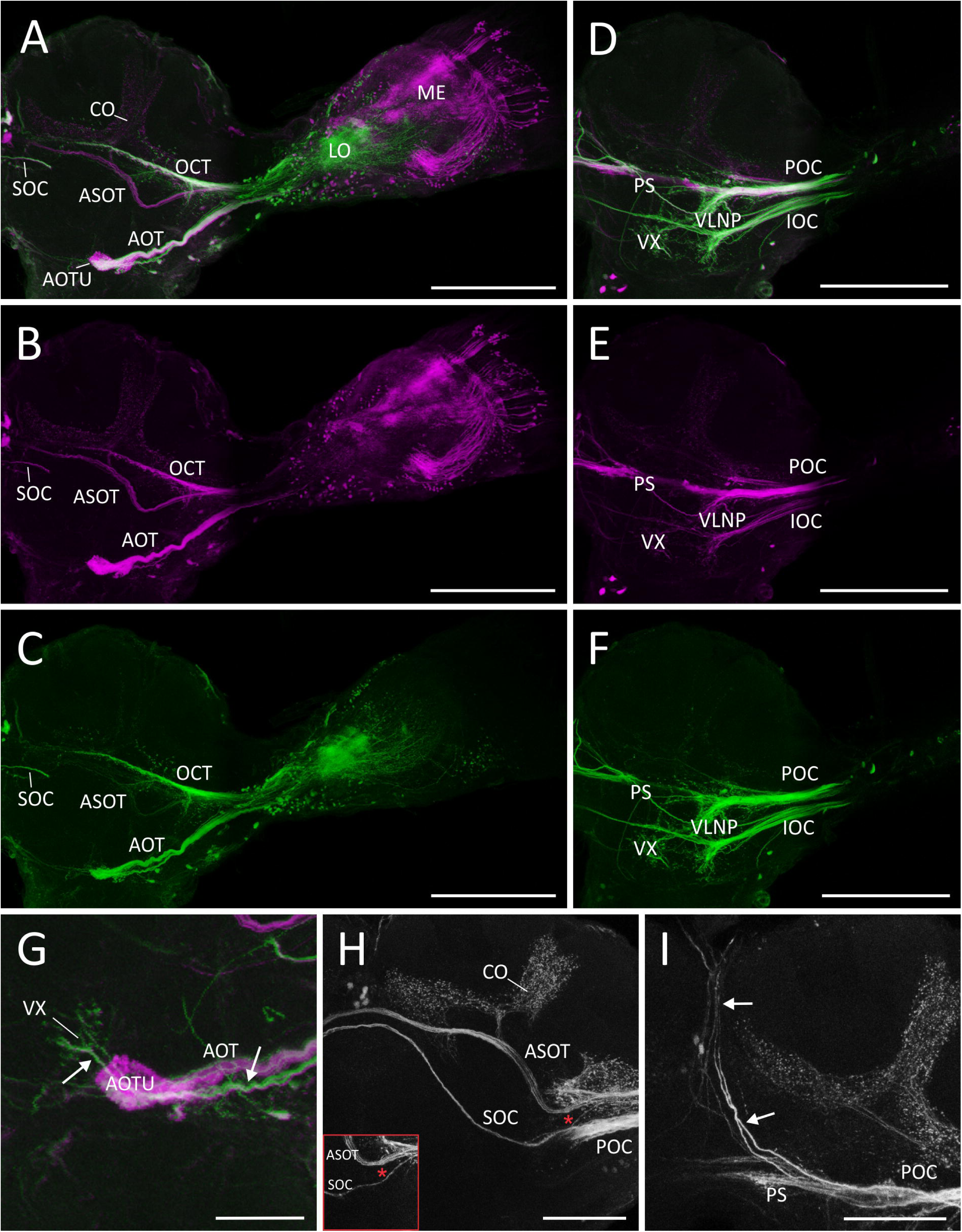
Visual projections from the optic lobes into the central brain. Anterograde staining obtained from dye injections into the medulla (ME; microruby in magenta or gray) and lobula (LO; Alexa 488 dextran in green). **A-C:** Visual fiber bundles of the ME (A,B) and the LO (A,C) located in the anterior brain. Anterior optic tract (*AOT*), anterior superior optic tract (*ASOT*), optical calycal tract (*OCT*) and superior optic commissure (*SOC*) comprise all projection neurons from ME and LO. The *ASOT* and the *OCT* project into the collar (CO) of the mushroom body and the *AOT* into the anterior optic tubercle (AOTU). Z-projection from a stack of 20 images with 5 µm step size. Scale bars = 200 µm. **D-F:** Visual fiber bundles of the ME (D,E) and LO (D,F) located in the posterior brain. The inferior (*IOC*) and posterior (*POC*) optic commissure comprise both neurons from ME and LO and project into the ventrolateral neuropils (VLNP) and the ventral complex (VX). In addition, the *POC* exhibits projections into the posterior slope (PS). Z-projection from a stack of 14 images with 5 µm step size. Scale bars = 200 µm. **G:** The *AOT* is accompanied by some neurons, which run into deeper layers of the brain and innervate there the ventral complex (neurons are indicated by white arrows). Z-projection from a stack of 24 images with 5 µm step size. Scale bar = 50 µm. **H:** The *SOC* exhibits no projections into the protocerebrum. The *SOC* neurons originate either from the *POC* or ventral to the *ASOT* (indicated by the asterisk, inset). Z-projection from a stack of 13 images or eight images (inset) with 5 µm step size. Scale bar = 100 µm. **I:** The *POC* is accompanied by some neurons with projections to the ocellar tracts (indicated by white arrows). Z-projection from a stack of ten images with 5 µm step size. Scale bar = 100 µm.

**Figure 13:**
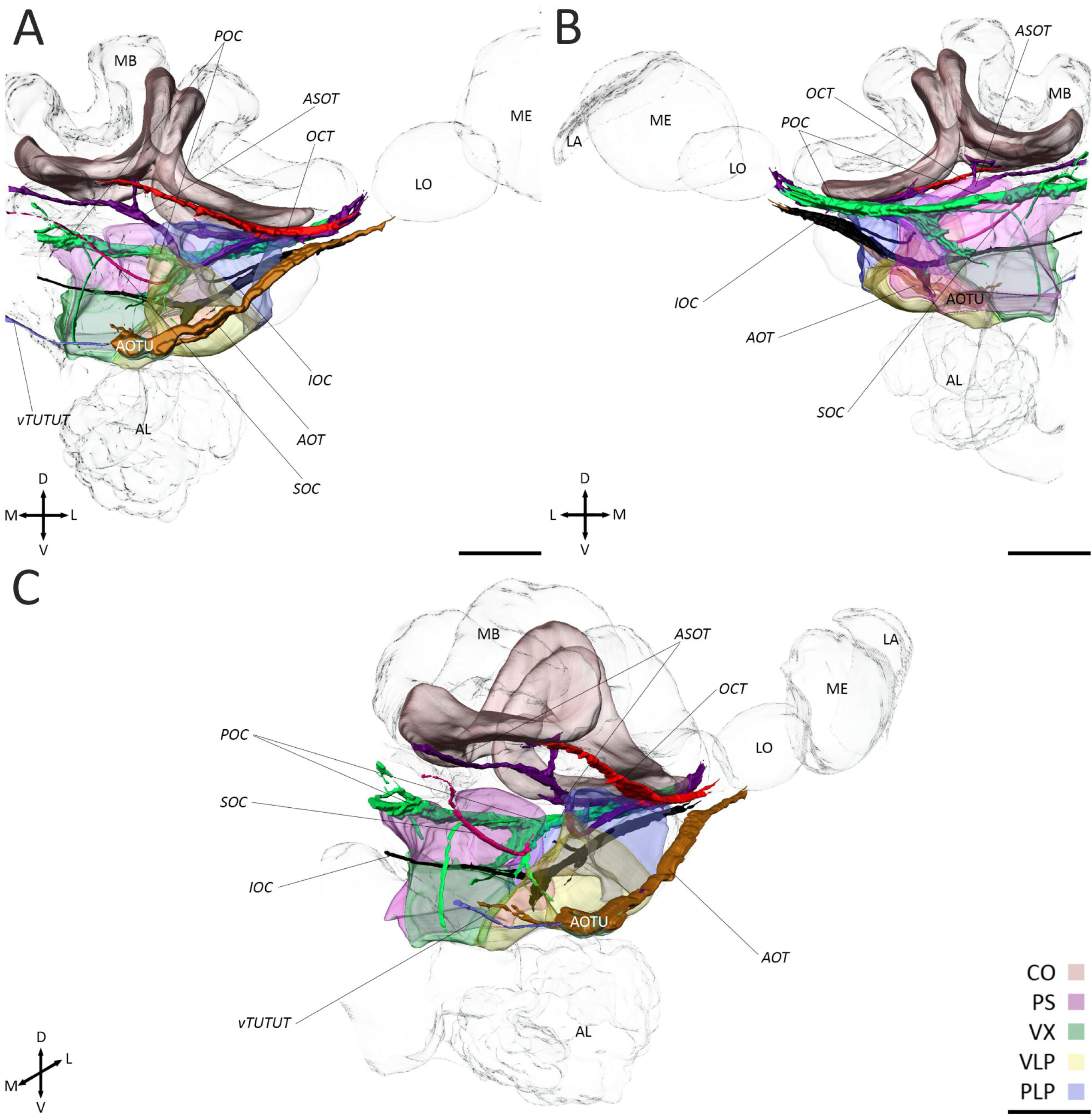
Surface reconstructions of the visual fiber bundles and their target neuropils. The surface reconstructions of the fiber bundles are based on anterograde staining and, in case of the neuropils on anti-synapsin labeling. Two major optic tracts (*AOT*: anterior optic tract; *OCT*: optical calycal tract) and four optic commissures (*ASOT*: anterior superior optic tract; *IOC*: inferior optic commissure; *POC*: posterior optic commissure; *SOC*: serpentine optic commissure) were found. The *OCT* projects into the collar (CO) of the ipsilateral mushroom body and the *AOT* into the anterior optic tubercle (AOTU). The AOTUs are interconnected by the ventral tubercle-tubercle tract (*vTUTUT*). In addition, the *AOT* is accompanied by some neurons which project into the ventral complex (VX). The *SOC* is the only commissure without any ramifications into the cerebrum. The *ASOT* exhibits arborizations into the CO, the *IOC* into the posterolateral protocerebrum (PLP), the ventrolateral protocerebrum (VLP) and the ventral complex (VX) and the *POC* into the PLP, the VLP, the VX and the posterior slope (PS). A: anterior view. B: posterior view. C: anteromedial oblique view. LA: lamina; LO: lobula; ME: medulla. Scale bars = 100 µm.

**Figure 14:**
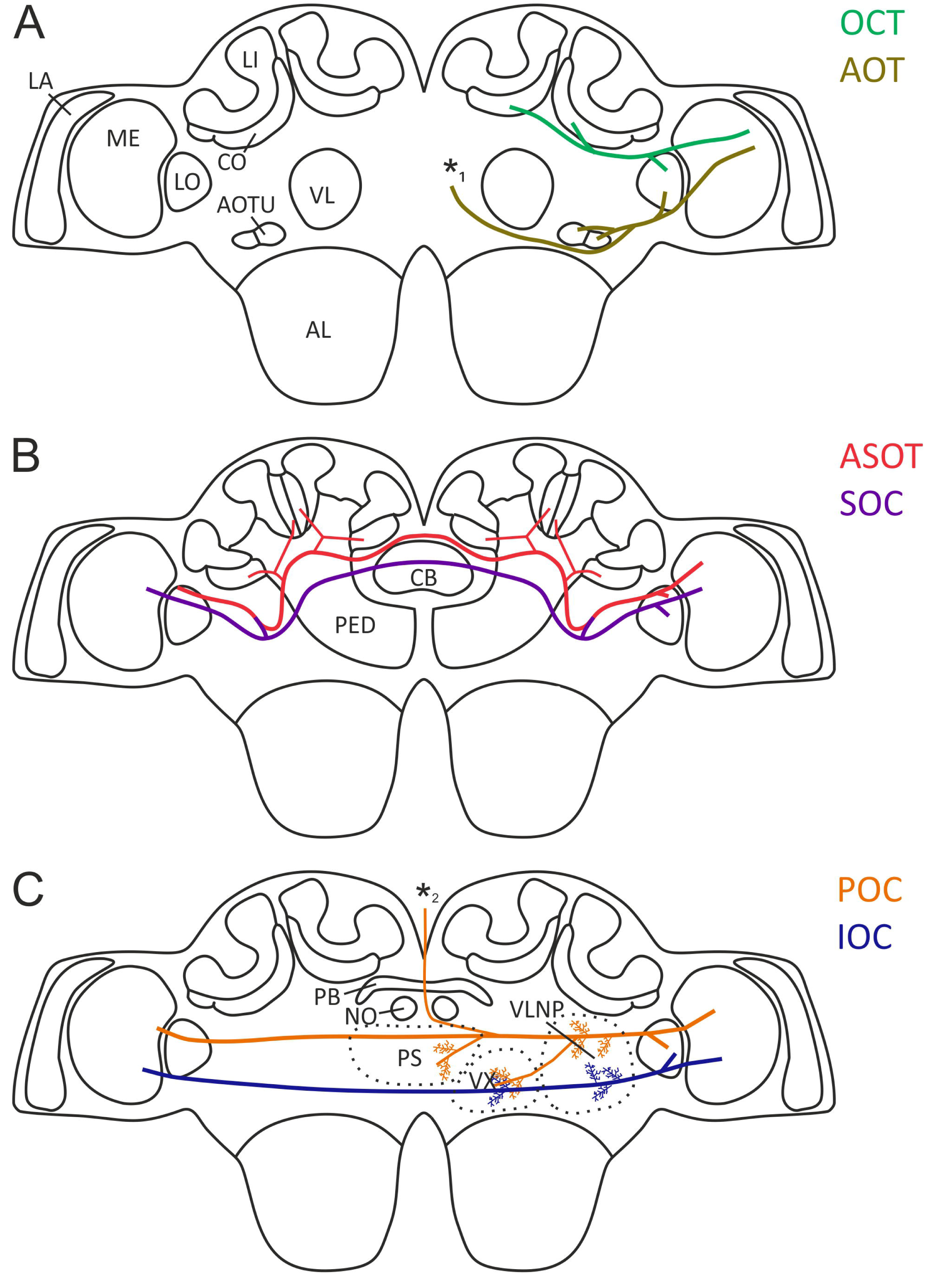
Schematic drawing of the visual fiber bundles of the *Cataglyphis nodus* brain. The overview of the optical tracts and commissures based on anterograde staining received from dye injections into the medulla (ME; microruby) and lobula (LO; Alexa 488 dextran). **A:** The optical calycal tract (*OCT*) and the anterior optic tract (*AOT*) are situated in the anterior part of the brain. The *OCT* projects into the ipsilateral collars (CO) of the mushroom bodies (MB) and the *AOT* into the ipsilateral anterior optic tubercle (AOTU). Both tracts receive input from the medulla (ME) and lobula (LO). The *AOT* is additionally accompanied by some neurons, which run into deeper layers of the brain, innervating the ventral complex (VX, indicated by asterisk 1). **B:** The anterior superior optic tract (*ASOT*) and the serpentine optic commissure (*SOC*) are situated more posteriorly than *AOT* and *OCT* but still relatively far anterior in the brain. Both commissures connect the ME and LO with their contralateral counterparts. The *ASOT* has additional ramifications into the ipsi- and contralateral COs. **C:** The posterior optic commissure (*POC*) and the inferior optic commissure (*IOC*) are situated in the posterior part of the brain. Both commissures connect the MEs and LOs between the hemispheres. The *POC* has ramifications into the dorsal region of the ventrolateral neuropils (VLNP), the ventral complex (VX) and the posterior slope (PS). In addition, some neurons, which accompany the *POC* connect the optic lobes with the ocelli (indicated by asterisk 2) The *IOC* innervates the ventrolateral part of the VLNP and the VX. AL: antennal lobe; CB: central body; LA: lamina; LI: lip; NO: noduli; PB: protocerebral bridge; PED: pedunculus.

In addition to these two fiber bundles, we found further tracts and commissures that project into different regions of the central brain or interconnect the OLs. We identified a very thin and serpentine shaped commissure - the serpentine optic commissure (*SOC*) (Figure 12h), which has also been described in the honey bee (Hertel and Maronde 1987; Hertel et al. 1987). The general shape of this extremely thin commissure has strong similarities to the *ASOT*. In contrast to the *ASOT*, the *SOC* is situated slightly more posteriorly in the brain and consists of fibers from only a few neurons, which project from the ME and LO to their bilateral counterparts without any further arborizations into other neuropils (Figure 12a-c, h, 13, 14b). The *SOC* neurons enter the central brain either ventrally to the *ASOT* (anterior in the brain) or more posteriorly, together with the neurons of the posterior optic commissure (*POC*) (Figure 12h). All *SOC* neurons merge at the inferolateral edge of the CL before they together pass the PE through the CL and cross the midline anterior to the CBU slightly ventral to the *ASOT* (Figure 12h, 14b).

Another very prominent tract in all insect brains investigated so far is the anterior optic tract (*AOT*, Figure 12a-c, g, 13, 14a). In *Cataglyphis*, the *AOT* is situated inferolaterally of the SLP and anteriorly to the LH. Among others, it transfers information from the sky polarization pattern from the optic lobes to the AOTU (el Jundi et al. 2011; Pfeiffer and Kinoshita 2012; Zeller et al. 2015). In *Cataglyphis*, the *AOT* is accompanied by additional neurons that pass the AOTU and run into the VX (Figure 12g, 13, 14a). Both subunits of the AOTU as well as the VX are supplied by projection neurons of the ME and the LO (Figure 12a-c, g).

In addition, we found two further commissures, the posterior optic commissure (*POC*) and the inferior optic commissure (*IOC*). Both commissures have previously been reported in honey bees (DeVoe et al. 1982; Mobbs 1984; Hertel and Maronde 1987; Hertel et al. 1987) and in the ant *Camponotus rufipes* (Yilmaz et al. 2016). In *Cataglyphis*, *POC* and *IOC* contain projection neurons of the ME and the LO (Figure 12d-f). The *POC* is the thickest visual fiber bundle in the *Cataglyphis* brain and easily recognizable in the immunostaining, even without any further anterograde tracing. After the commissure emerges from the optic lobes, it enters the cerebrum just ventral to the posterior part of the lateral calyx and exhibits many ramifications into the VLNPs (Figure 12d-f, 13, 14c). From there, the *POC* runs inferior to the calyces and superior to the PLP before it bifurcates into two relatively thick branches. One neuronal bundle exits the commissure ventrally and innervates the ventromedial neuropils (Figure 12d-f, 13, 14c). The remaining neurons of the *POC* continue along a more or less straight line in between the CA and the PS before they cross the midline above the IB (Figure 12d-f). Close to the midline, some of these neurons bifurcate into the ocellar tracts (Figure 12i). The *IOC* lies inferior to the *POC* (Figure 12d-f, 13, 14 c). In *Cataglyphis*, this commissure separates PLP (superior) and VLP (inferior) as well as CL/CRE (superior) and VX (inferior) before it crosses the midline of the brain. Lateral to the *l-ALT*, many neurons of the *IOC* leave the commissure ventrally and innervate the VLNPs. In addition, the *IOC* gives rise to medial arborizations into the VX (Figure 12d-f, 13, 14 c).

## Discussion

In this study, we examined the brain of the worker caste in the thermophilic ant *Cataglyphis nodus*, a favorable experimental model for the study of long-distance navigation. These ants are perfectly adapted to harsh environments with very high ground temperatures and scattered food resources, which they predominantly find during largely visually-guided foraging trips. Overall, we reconstructed 25 paired and 8 unpaired synapse-rich neuropils, defined 30 fiber tracts, among them 6 fiber tracts that provide new insights into the complexity of the visual system of the ants. A comparably detailed description of an insect brain that also includes the brain regions of the CANP currently exists only for the fruit fly *Drosophila melanogaster* (Pereanu et al. 2010; Ito et al. 2014), the monarch butterfly *Danaus plexippus* (Heinze and Reppert 2012), the ant *Cardiocondyla obscurior* (Bressan et al. 2015), the dung beetle *Scarabaeus lamarcki* (Immonen et al. 2017) and the desert locust *Schistocerca gregaria* (von Hadeln et al. 2018). The orientation, structure and overall layout of the *Cataglyphis* brain show large similarities with the honey bee brain (Brandt et al. 2005; Ribi et al. 2008). This is not surprising, as the brains of hymenopterans exhibit a similar general layout, including large and complex mushroom bodies. A most striking difference is the orientation of the brain. While the central brains of other insects, including dipterans, lepidopterans and coleopterans (el Jundi et al. 2009; Heinze and Reppert 2012; Ito et al. 2014; Immonen et al. 2017) tilt during the metamorphic development by 90° (Huetteroth et al. 2010), the brains of hymenopterans do not undergo such a modification. Thus, the central brain of e.g. ants, honey bees, wasps, and also hemimetabolous insects like locusts (Brandt et al. 2005; Kurylas et al. 2008; Groothuis et al. 2019) are oriented perpendicular compared to e.g. the fly brain. It was therefore not trivial to compare and identify similar brain regions in the ant brain using those described in the fly brain. Nevertheless, our description of the neuropils, tracts, fibers and commissures can serve as a reliable base for the brains of other ant species, bees, and wasps.

### Mushroom bodies and visual input

*Cataglyphis* ants, like honey bees, possess very prominent and complex mushroom bodies (Figure 1, 3): each MB consists of a medial and a lateral calyx, and each of these calyces, in turn, can further be subdivided into a collar and lip region. This subdivision most likely is a consequence of the high importance of visual memory, in addition to olfactory memory (Mobbs 1984; Hertel and Maronde 1987; Gronenberg 1999; Paulk and Gronenberg 2008; Yilmaz et al. 2016). Therefore, the MBs in *Cataglyphis* represent a truly multimodal integration center (Figure 6a, b, 12a-c, h) (Gronenberg 1999, 2001). The expansion of visual innervation of the MB calyx in species along the hymenopteran lineage first occurs in species with parasitoid lifestyles as well as in all social species, and, therefore, was interpreted as a trait that co-occurred with advanced spatial orientation (Farris and Schulmeister 2010). The importance of the MBs for learning and memory has been extensively demonstrated in honey bees (e.g. Erber et al. 1980; Menzel 1999, 2001; Szyszka et al. 2008) and can be assumed to serve a similar function in *Cataglyphis*. Although the exact number of Kenyon Cells (KCs) of the MBs in *Cataglyphis* is not known, it can be assumed to be very high compared to many other insects. For example, the MB of the ant *Camponotus rufipes* houses around 130,000 KC (Ehmer and Gronenberg 2004) and the MB collar in *Cataglyphis* contains an estimated number of 400,000 microglomeruli (Grob et al. 2017; Grob et al. 2019). This suggest that KC numbers in *Cataglyphis* MBs are at least as high as in *Camponotus* and a decent number is associated with the large collar region (Figure 3) (Gronenberg and Hölldobler 1999). In general, the number of KCs lies in a comparable magnitude in the honey bees MBs (340,000 KC) (Witthöft 1967; Strausfeld 2002; Rössler and Groh 2012), which is substantially higher than 2,500 KCs that were counted in *Drosophila* MBs (Fahrbach 2006). The discrepancy between *Drosophila* and hymenopteran MBs most likely concerns differences in navigational abilities and the socio-ecological context of social Hymenoptera. As outlined above, the correlation between advanced spatial orientation with the visual input and expansion of the MBs in higher Hymenoptera was proposed in several studies (Farris and Schulmeister 2010; Farris 2013, 2015).

Our study shows that several fibers and commissures transfer visual input into the MBs of *Cataglyphis nodus,* which likely causes the relatively large collar region containing a high number of input synaptic complexes (microglomeruli) (Stieb et al. 2012; Grob et al. 2017). The large MBs, in general, and the substantial amount of visual input into the collar are in line with the excellent abilities of *Cataglyphis nodus* and other *Cataglyphis* species to form visual memories for landmarks and/or panoramic sceneries as shown or highlighted more recently (Fleischmann et al. 2016; Fleischmann et al. 2018; for reviews see: Rössler 2019; Zeil and Fleischmann 2019). *Cataglyphis* foragers have to learn the visual features of their nest surroundings in order to find their way to their nest entrance when homing back from foraging trips (Wehner and Menzel 1969; Wehner and Räber 1979; Cruse and Wehner 2011). Several studies suggest that the latter is achieved by taking snapshots of the panorama during specific learning walks when the ants leave their nest for the first time (Müller and Wehner 1988; Fleischmann et al. 2017). This visual information, most likely, is stored in the vast number of associative visual microcircuits that involve many thousands of Kenyon cells. It appears counterintuitive, however, that the basal ring, a substructure known from honey bees, is not a distinct structure in *Cataglyphis* MBs, especially as the basal ring is known as a multimodal sensory input region integrating visual and olfactory stimuli (Gronenberg 1999, 2001). However, also in other ant species, the basal ring is largely reduced, possibly absent, or indistinguishable from the collar (Gronenberg 1999; Ehmer and Gronenberg 2004). The reasons for this remain unclear.

### Antennal lobes and dual olfactory pathway

Recent studies showed that the ALs of *C. nodus* worker contain about 226 glomeruli, which is in a similar range to other *Cataglyphis* species such as *C. fortis* (∼ 198) and *C. bicolor* (∼ 249) (Stieb et al. 2011). However, this number is substantially smaller compared to high glomeruli numbers in other ant species, such as *Camponotus floridanus* (∼ 434) (Zube et al. 2008), *Camponotus japonicus* (∼ 438) (Nishikawa et al. 2008), the wood ant *Formica rufibarbis* (∼ 373) (Stieb et al. 2011), or the high numbers in leaf cutting ants, for example *Atta vollenweideri* (Kelber et al. 2009). The reduced number of glomeruli might reflect the different ecology of *Cataglyphis* ants that, compared to all other species mentioned above, primarily rely on visual information instead of pheromonal cues during foraging (Ruano et al. 2000; reviewed by: Wehner 2003). On the other hand, studies have shown that *Cataglyphis* do use olfactory cues during cross wind orientation while searching for food (Steck et al. 2011; Buehlmann et al. 2014). However, both the role of pheromones and food odors may be minor compared to ants with reduced visual capabilities. In any case, compared to non-social insects, e.g. *Drosophila* (∼ 40) (Stocker et al. 1990; Laissue et al. 1999; Wong et al. 2002), lepidopterans (∼ 50-70) (e.g. Masante-Roca et al. 2005; Varela et al. 2009; Montgomery and Ott 2015) and dung beetles (∼ 85) (Immonen et al. 2017), the number of olfactory glomeruli in *Cataglyphis* still are comparatively high most likely due to social interactions and communication inside the nest using cuticular hydrocarbon cues (Hölldobler and Wilson 1990).

We also found that *Cataglyphis* possess a dual olfactory pathway, similar to the previously described feature in various other higher hymenopterans (Kirschner et al. 2006; Zube et al. 2008; Rössler and Zube 2011; Brill et al. 2015; Couto et al. 2016). Like in other Hymenoptera, projection neurons of different subpopulations of glomeruli and projection neurons form the major tracts *m-ALT*, *l-ALT* and the three thinner *ml-ALTs* in *Cataglyphis* (Figure 6a). This feature appears more complex compared to most other insect species, with a smaller number of antennal lobe output tracts (reviewed by: Galizia and Rössler 2010). Functional studies suggest that both parallel processing and coincidence coding are employed within this system leading to enhanced coding capabilities for these insects with highly complex olfactory environments and behaviors (Müller et al. 2002; Brill et al. 2013; Rössler and Brill 2013; Brill et al. 2015; Carcaud et al. 2015).

### Optic lobes

The OLs in the *Cataglyphis* brain are relatively large in comparison to other ant species (Figure 1) (Gronenberg and Hölldobler 1999). Enlarged OLs are otherwise typically present in winged, reproductive castes or visually hunting ants, which underpins the high relevance of visual stimuli for visually based navigation in *Cataglyphis* workers. The OLs of *Cataglyphis* show the typical structure with lamina, medulla and lobula (Figure 2) (Gronenberg and Hölldobler 1999; Brandt et al. 2005; Gronenberg 2008). In contrast to Diptera, Coleoptera, Lepidoptera or Tricoptera, Hymenoptera possess one coherent lobula (Dettner and Peters 2011) instead of a lobula complex that consists of two distinct areas: lobula and lobula plate. In many insects, the lobula plate is a center for motion vision that encodes optic-flow information (Hausen 1976; Hausen 1984; Joesch et al. 2008). However, studies in *Cataglyphis bicolor* have shown that the ants perceive optic-flow information and use it for distance estimations (Pfeffer and Wittlinger 2016). This raises the question how optic flow information is processed in the ant brain (and in general in Hymenoptera), more explicitly whether the lobula can further be subdivided into different functional subregions, similar to what has been shown for the praying mantis and the locust (Kurylas et al. 2008; Rosner et al. 2017).

### Central complex, anterior optic tubercle and lateral complex

In *Cataglyphis*, the anterior optic tubercle (AOTU) comprises an upper and a lower subunit which seems to be highly conserved in all insects studied so far (Strausfeld and Okamura 2007; el Jundi et al. 2010; Mota et al. 2011; Heinze and Reppert 2012; Montgomery et al. 2016; Immonen et al. 2017; von Hadeln et al. 2018). In all insects, sky compass neurons project into the lower subunit of the AOTU whereas the upper division is associated with chromatic, unpolarized light vision (Mota et al. 2011; Pfeiffer and Kinoshita 2012; Mota et al. 2013; Zeller et al. 2015). In *Cataglyphis*, the AOTUs on both sides are connected via the tubercle-bulb tract (*TUBUT*, Figure 9c) and transmit sky compass information to the bulb, similar to what has been shown in other insects studied so far (Pfeiffer et al. 2005; el Jundi and Homberg 2010; Heinze and Reppert 2012).

While the boundaries of the lateral accessory lobe (LAL) of the lateral complex (LX), a region that mediates motor output to the ventral nerve cord (Namiki and Kanzaki 2016; Turner-Evans and Jayaraman 2016), are more difficult to align, the bulbs are easily recognizable in *Cataglyphis* due to the existence of large microglomerular synaptic structures (Figure 3g, 4d). These microglomeruli gate sky compass and other visual information before entering the lower unit of the central body (CBL) via tangential neurons (TN) (Pfeiffer and Homberg 2007; Heinze and Reppert 2012; Seelig and Jayaraman 2013). Interestingly, precocious stimulation of the ants with UV-light was shown to cause an increase in the number of microglomeruli in the bulbs of *Cataglyphis fortis* indicating a substantial level of structural plasticity in these synaptic circuits (Schmitt et al. 2016b). *Cataglyphis* has only one bulb per hemisphere, like *Drosophila* (Seelig and Jayaraman 2013), monarch butterflies (Heinze et al. 2013), and dung beetles (el Jundi et al. 2018) while locusts and honey bees possess two distinct bulbs or groups of microglomerular complexes (Heinze and Homberg 2008; Träger et al. 2008; el Jundi et al. 2014; Mota et al. 2016; von Hadeln et al. 2018). The reason for these differences are currently not known.

We could also define the isthmus tract in *Cataglyphis*. Via this tract, amongst others, visual information from the bulbs of the LX is transferred to the central complex (CX) as has been demonstrated in various insects (Homberg et al. 2011; el Jundi et al. 2014; Schmitt et al. 2016b; Sun et al. 2017). The overall layout of the CX in *Cataglyphis* (Figure 1, 4) appears similar to other insect species such as dung beetles (Immonen et al. 2017; el Jundi et al. 2018), bees (Brandt et al. 2005; Pfeiffer and Kinoshita 2012), *Drosophila* (Pereanu et al. 2010; Ito et al. 2014), monarch butterflies (Heinze et al. 2013) and locusts (Heinze et al. 2009; von Hadeln et al. 2018). Future studies using additional markers and single neuron labeling are needed to reveal the fine structure of the different components of the CX. However, the prominent input from the sky-compass system and similarities in general layout are highly suggestive for an important role of the CX in path integration in *Cataglyphis* ants, especially in the light of recent studies on the neuronal network that encodes the current and desired directions in the CX network (Seelig and Jayaraman 2015; Stone et al. 2017; Green et al. 2019).

### Central adjoining neuropils

Even though the function of most areas within the CANP is largely unknown, the use of genetic tools in *Drosophila* promoted functional studies of the neuronal circuits in this brain region. For instance, novel calcium imaging tools in behaving flies could demonstrate the participation of different neuropils of the CANP in walking behavior (Aimon et al. 2019). To better relate and compare studies in other insects with *Cataglyphis,* we subdivided the CANP into subunits using glial boundaries, fiber tracts, f-actin phalloidin staining as well as synapsin-ir and 5-HT-ir as criteria for defining borders. This first map of the CANPs will allow us to study these brain regions in more detail in the future, particularly for investigating the distribution of neurotransmitters, neuromodulators, individual neurons and circuits, or gene expression profiles and compare these features directly with the situation in other insects.

Most of the described neuropils within the CANP of *Drosophila* do also exist in the brain of *Cataglyphis* (Figure 7, 8). Since the enormous MBs of *Cataglyphis* occupy large parts of the central brain, the location and the size of the CANP components differ in some cases compared to the CANP in *Drosophila* and other insects investigated for this region so far. For instance, the crepine encases the complete medial lobe of the MB in *Drosophila* (Ito et al. 2014) but appears much smaller and only around the tip of the medial lobe in *Cataglyphis* (Figure 7h, i). Nevertheless, the general layout and location of the CANP components in *Cataglyphis* are very similar to other insects and appear as diverse as in dung beetles, fruit flies, locusts and monarch butterflies (Heinze and Reppert 2012; Ito et al. 2014; Immonen et al. 2017; von Hadeln et al. 2018). However, due to the lack of clear landmarks, we did not further subdivide some brain areas within the CANP of *Cataglyphis*. This concerns the clamp (CL), the posterior slope (PS), the ventrolateral complex (VX) and the ventrolateral protocerebrum (VLP). Using the present template as a basis, the use of more diverse molecular markers might open up new insight into subdivisions in the future.

Our anterograde staining highlighted some subdivisions of the CANP in *Cataglyphis*, which largely correspond with previous investigations in other Hymenoptera (Zube et al. 2008; reviewed by: Galizia and Rössler 2010; Rössler and Zube 2011) and *Drosophila* (Tanaka et al. 2012a; Tanaka et al. 2012b). Thus, the LH, the superior intermediate protocerebrum (referred as “ring neuropil” in previous studies) and the ventrolateral neuropils receive olfactory information from the antennal lobes (Figure 6). In addition, the ventrolateral neuropils and the ventromedial neuropils receive visual input from the optic lobes (Figure 12a-g, 13, 14), which is consistent with data from honey bees (Hertel and Maronde 1987; Milde 1988; Maronde 1991), bumblebees (Paulk et al. 2008) and *Drosophila* (Strausfeld 1976; Otsuna and Ito 2006; Panser et al. 2016; Wu et al. 2016; Namiki et al. 2018). In flies, both brain regions are associated with the detection of directed motion and looming of objects (Ibbotson et al. 1991; Wicklein and Strausfeld 2000; Okamura and Strausfeld 2007; Wu et al. 2016; Klapoetke et al. 2017). These very conserved findings across species illustrate the importance and behavioral relevance of distinct CANP structures as high order integration centers in the insect brain.

### Visual projections in *Cataglyphis*

We confirmed two major optical tracts and four optic commissures in *Cataglyphis*. The anteriormost tract is the anterior optic tract (*AOT*, Figure 12a-c, 13, 14a). The *AOT* seems to be a most conserved optic tract in insects. It has been described in diverse insect orders such as Blattodea (Reischig and Stengl 2002; Rosner et al. 2017), Coleoptera (Immonen et al. 2017), Diptera (Power 1943; Strausfeld 1976; Fischbach and Lyly-Hünerberg 1983; Omoto et al. 2017; Adden et al. 2019), Lepidoptera (Strausfeld and Blest 1970; Collett 1972), Orthoptera (Homberg et al. 2003; Pfeiffer et al. 2005) and Hymenoptera including different *Cataglyphis* species (Mota et al. 2011; Pfeiffer and Kinoshita 2012; Held et al. 2016; Schmitt et al. 2016b; Grob et al. 2017). The *AOT* relays optic motion information (Collett 1972; DeVoe et al. 1982; Paulk et al. 2008) as well as chromatic and polarization cues (Pfeiffer et al. 2005; Kinoshita et al. 2007; Mota et al. 2011) to the AOTU. In *Cataglyphis*, the *AOT* consists of projection neurons from the medulla and lobula (Figure 12a-c). Surprisingly, we also found that a few isolated neurons accompany the *AOT* but arborize into the VX instead of the AOTU (Figure 12g). Whether these neurons transmit similar information or completely different visual properties than other *AOT*-neurons requires future functional studies.

The optical calycal tract (*OCT*) exclusively projects into the ipsilateral collars in the *Cataglyphis* brain (Figure 12a-c, 13, 14a). To our knowledge, this tract has not been described before in any insect. Three different optic tracts, the anterior superior optic tract (*ASOT*), the anterior inferior optic tract (*AIOT*) and the lobula tract (*LOT*) are known in honey bees (Mobbs 1984; Ehmer and Gronenberg 2002). In *Cataglyphis*, we did not find any evidence for a distinct *AIOT* or *LOT*. Instead, a very small subset of lobula neurons join the *ASOT* formed by dorsal and ventral medullar neurons (Grob et al. 2017) and project into the calyces of both hemispheres (Figure 14b). The *ASOT* has exclusively been described in Hymenoptera, whereas other insects possess another anterior optical commissure that we could not find in *Cataglyphis* – the great commissure (*GC*) (Strausfeld 1976; Ito et al. 2014; Immonen et al. 2017; von Hadeln et al. 2018). Whether the *GC* is homologous to the *ASOT*, meaning that the *ASOT* is a variation of the *GC,* is unknown. One of the major differences between both commissures is that the *GC* completely lacks any projections into the calyces, whereas the *ASOT* has prominent projections into the lateral and medial calyces (Strausfeld 1976; Farris 2005; Ito et al. 2014).

We also identified a commissure that exclusively interconnects the OLs in the *Cataglyphis* brain - the serpentine optic commissure (*SOC*; Figure 12a-c, h, 13, 14b), which has previously been described in the brains of the cockroach *Leucophaea maderae* (Loesel and Homberg 2001; Reischig and Stengl 2002), crickets (Honegger and Schürmann 1975; Tomioka et al. 1994) and the honey bee (Hertel and Maronde 1987). In all of these insects, the *SOC* comprises only very few neurons originating from the lobula and medulla, which is consistent with our findings in *Cataglyphis* (Figure 12a-c). These neurons might respond to moving objects and play a role in tracking of objects as it has been shown in other insects (Hertel and Maronde 1987; Loesel and Homberg 2001).

The posterior optic commissure (*POC*) and the inferior optic commissure (*IOC*) are situated more posteriorly in the *Cataglyphis* brain. Here, the *POC* is a very prominent commissure. It was described in numerous insect species including Hymenoptera (e.g. Honegger and Schürmann 1975; Hertel and Maronde 1987; Reischig and Stengl 2002; Ito et al. 2014; Yilmaz et al. 2016; Immonen et al. 2017; von Hadeln et al. 2018). In contrast, the *IOC* has only been described in few insect species such as the cockroach *Leucophaea maderae* (Reischig and Stengl 2002), crickets (Honegger and Schürmann 1975) and Hymenoptera (Mobbs 1984; Hertel and Maronde 1987; Paulk et al. 2008; Yilmaz et al. 2016). These tracts transfer, amongst others, information from the OL to the VMNPs and the VLNPs. Electrophysiological recordings in bees showed that these neurons, for the most part, are achromatic (Hertel and Maronde 1987; Paulk et al. 2009). Further results associated the neurons of the *POC* with the localization of stationary targets, while the *IOC* neurons respond to directional movements of objects (Hertel and Maronde 1987; Maronde 1991).

## Acknowledgements

We would like to thank the Greek government and the management boards of the Schinias and Strofylia National Park for the permission to excavate and transfer the *Cataglyphis* ants to Germany. We especially thank Maria Trivourea, Vasiliki Orfanou and Georgia Karamperou for the warm welcome and their support during our field work. We are very grateful to Christos Georgiadis for his longstanding cooperation administrative help, and for guiding us to the *Cataglyphis* nests. We also thank the field assistants who defied the heat in the field and helped to excavate the *Cataglyphis* nests. Special thanks go to Erich Buchner and Christian Wegener for kindly providing the anti-synapsin antibody.

## Role of the authors

Study concept and design: WR, BeJ, JH, EA. Preparation and acquisition of data: EA, KG, JH. Analyses and interpretation of data: JH, EA, KG, BeJ, WR. Drafting of the manuscript: JH. Critical review of the manuscript: EA, BeJ, WR. Obtained funding: WR. All authors approved the final version of the manuscript for submission.

## Funding

The study was supported by the German Research Foundation (DFG), DFG Ro1177/7-1 and DFG equipment grant INST 93/829-1, both to WR.

## CONFLICT OF INTEREST STATEMENT

The authors declare no conflicts of interest.

## Abbreviations

5-HT: serotonin
AL: antennal lobes
AMMC: antennal mechanosensory and motor center
*AMMCC*: antennal mechanosensory and motor center commissure
AN: antennal nerve
*AOT*: anterior optic tract
AOTU: anterior optic tubercle
ATL: antler
*ASOT*: anterior superior optic tract
AVLP: anterior ventrolateral protocerebrum
BU: bulb
CA: calyx
CAN: cantle
CANP: central adjoining neuropil
CB: central body
CBL: central body lower division
CBU: central body upper division
CL: clamp
CO: collar
CRE: crepine
CRG: cerebral ganglia
CX: central complex
EB: ellipsoid body
FB: fan-shaped body
FLA: flange
GNG: gnathal ganglion
*hVLPF*: horizontal ventrolateral protocerebrum fascicle
IB: inferior bridge
*IFS*: inferior fiber system
INP: inferior neuropils
*IOC*: inferior optic commissure
*IT*: isthmus tract
KC: kenyon cells
LA: lamina
LAL: lateral accessory lobe
*LALC*: lateral accessory lobe commissure
*l-ALT*: lateral antennal lobe tract
LCA: lateral calyx
*LEF*: lateral equatorial fascicle
LI: lip
LH: lateral horn
LO: lobula
LX: lateral complex
*m-ALT*: medial antennal lobe tract
MB: mushroom body
*MBDL*: median bundle
MCA: medial calyx
ME: medulla
*MEF*: medial equatorial fascicle
ML: medial lobe
*ml-ALT*: mediolateral antennal lobe tract
*mTUTUT*: medial tubercle-tubercle tract
NGS: normal goat serum
NO: noduli
*OCT*: optical calycal tract
OL: optic lobes
ORN: olfactory receptor neuron
PB: protocerebral bridge
PBS: phosphate-buffered saline
*PCT*: protocerebral-calycal tract
PED: pedunculus
PENP: periesophageal neuropils
*PLF*: posterior lateral fascicle
PLP: posterolateral protocerebrum
*POC*: posterior optic commissure
PRW: prow
PS: posterior slope
PVLP: posterior ventrolateral protocerebrum
*PYF*: pyriform fascicle
SAD: saddle
*SEC/SAC*: superior ellipsoid/arch commissure
*SFS*: superior fiber system
SIP: superior intermediate protocerebrum
SLP: superior lateral protocerebrum
*SLPT*: anterior superior lateral protocerebrum tract
SMP: superior medial protocerebrum
SMPT: superior medial protocerebrum tract
SNP: superior neuropils
*SOC*: serpentine optic commissure
SPL: serpentine layer
*sPLPC*: superior posterolateral protocerebrum commissure
TN: tangential neurons
*TUBUT*: tubercle-bulb tract
VL: vertical lobe
VLNP: ventrolateral neuropils
VLP: ventrolateral protocerebrum
VMNP: ventromedial neuropils
*vTUTUT*: ventral tubercle-tubercle tract
VX: ventral complex.

